# Intra-species signaling between *Pseudomonas aeruginosa* genotypes increases production of quorum sensing controlled virulence factors

**DOI:** 10.1101/2020.04.29.068346

**Authors:** Dallas L. Mould, Nico J. Botelho, Deborah A. Hogan

## Abstract

The opportunistic pathogen *Pseudomonas aeruginosa* damages hosts through the production of diverse secreted products, many of which are regulated by quorum sensing. The *lasR* gene, which encodes a central quorum-sensing regulator, is frequently mutated, and loss of LasR function impairs the activity of downstream regulators RhlR and PqsR. We found that in diverse models, the presence of *P. aeruginosa* wild type causes LasR loss-of-function strains to hyperproduce RhlR/I-regulated antagonistic factors, and autoinducer production by the wild type is not required for this effect. We uncovered a reciprocal interaction between isogenic wild type and *lasR* mutant pairs wherein the iron-scavenging siderophore pyochelin, specifically produced by the *lasR* mutant, induces citrate release and cross-feeding from wild type. Citrate stimulates RhlR signaling and RhlI levels in LasR-but not in LasR+ strains, and the interactions occur in diverse media. Co-culture interactions between strains that differ by the function of a single transcription factor may explain worse outcomes associated with mixtures of LasR+ and LasR loss-of-function strains. More broadly, this report illustrates how interactions within a genotypically diverse population, similar to those that frequently develop in natural settings, can promote net virulence factor production.

## Introduction

Genetic diversity frequently arises and persists within clonally-derived bacterial and fungal populations in chronic infections and healthy microbiomes, and recent data highlight that this heterogeneity can pose challenges to clearance and treatment (1-3). Genotypic and phenotypic complexity has been particularly well-documented in the chronic lung infections associated with the genetic disease cystic fibrosis (CF), and these studies have convincingly demonstrated that within a species, a common set of genes is under selection across strains and hosts (4-10).

*P. aeruginosa* loss-of-function mutations in *lasR* (LasR-) are very commonly found in CF isolates, strains from acute infections, and from environmental sources (11-15). Although infection models often show that *lasR* loss-of-function mutants have reduced virulence compared to strains with functional LasR (LasR+) in several animal models (16, 17), the presence of LasR-strains is correlated with worse disease outcomes in acute and chronic infections (11, 12). There are several possible explanations for this apparent contradiction. Loss of lasR function confers some fitness advantages including altered catabolic profiles (18) and enhanced growth in low oxygen (19, 20). Further, some LasR-strains exhibit rewired regulation of quorum sensing (QS)-controlled exoproducts (21) and LasR-strains can activate QS signaling in response to products from other species (22) or in specific culture conditions (23, 24). LasR-strains are also frequently found among LasR+ *P. aeruginosa* strains where exoproducts can be shared or signal cross-feeding can occur (13).

*P. aeruginosa* LasR participates in the regulation of QS in conjunction with two transcription factors: RhlR and PqsR (MvfR). Each of these three regulators has one or more autoinducer ligands: 3-oxo-C12-homoserine lactone (3OC12HSL) for LasR, C4-homoserine lactone (C4HSL) for RhlR, and hydroxyalkylquinolones (Pseudomonas Quinolone Signal (PQS) and hydroxy-heptyl quinolone (HHQ)) for PqsR (25). In the regulatory networks described in the widely-used *P. aeruginosa* model strains, LasR is an upstream regulator of RhlR and PqsR signaling, and together these regulators control the expression of a suite of genes associated with virulence including redox-active small molecule phenazines (26-28), cyanide (29), proteases (30-33), and rhamnolipid surfactants important for surface motility, biofilm dispersal, and host cell damage (34-36).

Other traits that are often heterogeneous across *P. aeruginosa* isolates relate to strategies for iron acquisition. *P. aeruginosa* procures iron through the use of siderophores, including pyochelin (37-39) and pyoverdine (40), from heme or through a direct iron uptake system (41-43). It is common to encounter *P. aeruginosa* strains with loss of function mutations in genes encoding pyoverdine, the high affinity siderophore, but genes associated with use of the pyochelin siderophore, heme utilization, and the direct iron uptake are generally intact (44-46). Iron limitation can reduce the function of pathways that require abundant iron including the TCA cycle (47), and when iron access is low, *Pseudomonas spp.* release partially oxidized metabolic intermediates that accumulate at iron requiring steps (48, 49). Many other species are known to release partially-oxidized metabolic intermediates upon iron limitation (48, 50-52).

Here, we show that mixtures of *P. aeruginosa* LasR- and LasR+ strains have enhanced production of QS-controlled factors across medium types, culture conditions, and strain backgrounds, and that this is due to activation of RhlR likely through increased RhlI stability in LasR-strains. Our transcriptomic, genetic and biochemical studies led us to uncover a set of interactions in which Δ*lasR* production of the siderophore pyochelin was necessary for the activation of RhlR in Δ*lasR* when co-cultured with the LasR+ wild type (WT). Our studies suggested that Δ*lasR* responds to citrate in co-culture, and we found that pyochelin is sufficient to stimulate citrate release preferentially by WT cells. Citrate increased RhlI protein levels and RhlR activity in Δ*lasR* cells but not in the wild type. Together, these data highlight a set of complex, small molecule interactions between strains that differ by a single mutation and lead to increased production of exoproducts known to cause host damage.

## RESULTS

### *P. aeruginosa* Δ*lasR* over-produces pyocyanin in co-culture with wild type

We observed that mixtures of *P. aeruginosa* LasR+ and LasR-strains had high levels of total pyocyanin, a secreted, blue-pigmented phenazine. As shown in spot colony cultures of *P. aeruginosa* strain PA14 wild type (WT), Δ*lasR*, and WT / Δ*lasR* co-cultures, the strain mixture showed increased blue pigmentation (**Fig. 1A**) and a significant 2-fold induction of pyocyanin relative to either strain alone (**Fig. 1B**). Phenazine-deficient derivatives, Δ*phz* (Δ*phzA1-G1*Δ*phzA2-G2*) and Δ*lasRΔphz* (Δ*lasRΔphzC1C2*) were also included, and as expected, Δ*phz* and Δ*lasRΔphz*, showed no blue colony pigmentation (**Fig. 1A**) and no pyocyanin signal was evident above background levels (**Fig. 1B**). The higher levels of pyocyanin in WT / Δ*lasR* co-cultures relative to single-strain cultures was also observed on pH-buffered M63 medium and on artificial sputum medium indicating that the phenomenon occurred in diverse conditions (**Fig. S1A**). Co-cultures of clonally-derived LasR+ and LasR-clinical isolates collected from single respiratory sputum samples from chronically-infected individuals with cystic fibrosis also had increased production of a pyocyanin when LasR+ and LasR-strains were grown together relative to mono-culture levels (**Fig. S1B, C**).

**Figure 1.**
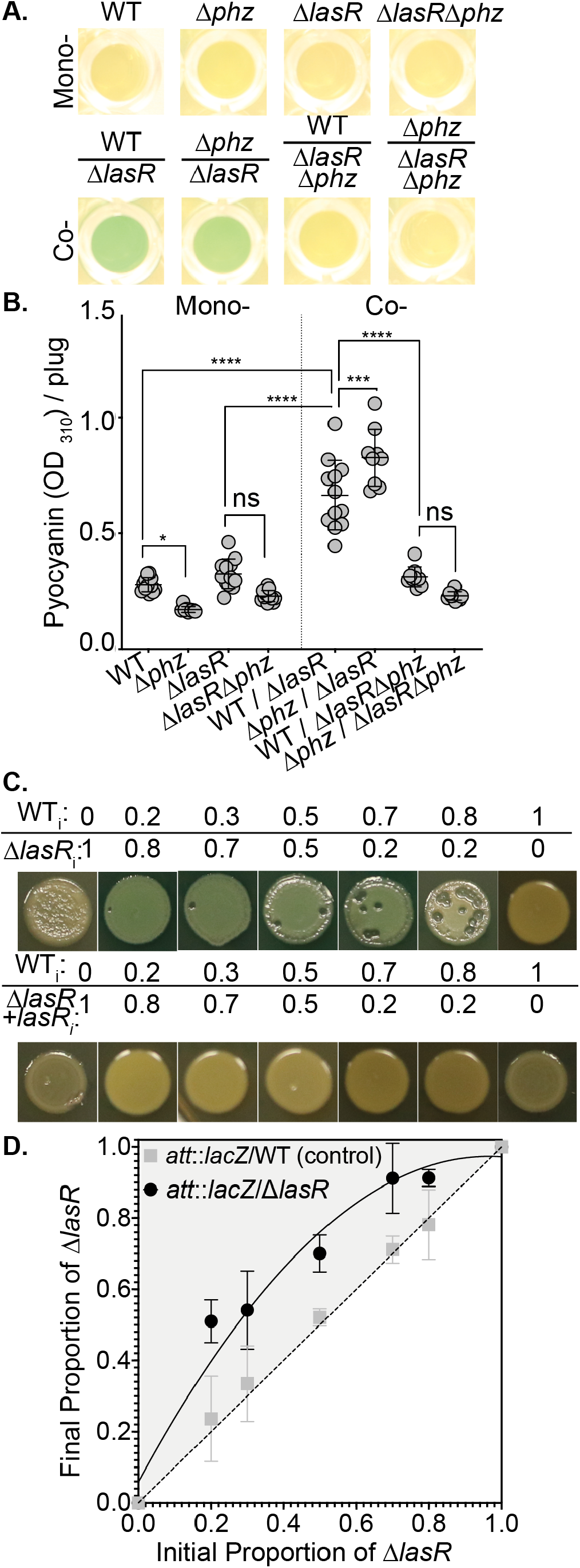
Δ*lasR* produces pyocyanin in wild type / Δ*lasR* co-cultures beyond monoculture concentrations. **A**. Wild type (WT), Δ*lasR*, and their phenazine-deficient derivatives (Δ*phz*) visualized from the bottom of the 96 well LB agar plate after 16 h growth as mono- (top) and co-cultures (bottom). **B**. Pyocyanin levels quantified for cultures described in A; ****, p<0.001. **C**. Pyocyanin production for wild type co-cultures with Δ*lasR* or Δ*lasR* complemented with the *lasR* gene at the native locus (Δ*lasR* + l*asR*) across several initial (_i_) proportions on LB for 20 h. **D**. Final proportions quantified after 16 h growth for WT and Δ*lasR* co-cultured with a WT tagged with *lacZ*. Experimental setup as described in **C**.

To assess individual strain contributions to increased pyocyanin in WT / Δ*lasR* co-cultures, we measured pyocyanin levels in co-cultures wherein one strain was replaced with its phenazine-deficient derivative. When Δ*lasR* was cultured with the phenazine biosynthesis mutant Δ*phz* (Δ*phz* / Δ*lasR*), we still observed increased blue pigmentation (**Fig. 1A**) and total pyocyanin levels were higher relative to monocultures (**Fig. 1B**). Surprisingly, pyocyanin levels were statistically higher in Δ*phz* / Δ*lasR* co-cultures relative to WT / Δ*lasR* (**Fig. 1B**). In contrast, WT / Δ*lasR*Δ*phz* co-cultures did not display the high pyocyanin phenotype, and pyocyanin concentrations in these co-cultures were more similar to those in WT monocultures (**Fig. 1A, B**). Pyocyanin levels in the WT / Δ*lasR*Δ*phz* co-cultures were below the limit of detection as there was no difference in pyocyanin levels when compared with Δ*phz* / *ΔlasRΔphz* co-cultures or Δ*phz* and Δ*lasRΔphz* monocultures. Collectively, these data suggested that WT / Δ*lasR* co-cultures produced more pyocyanin than either monoculture alone, and that pyocyanin was contributed by the Δ*lasR* strain.

The higher levels of pyocyanin in WT / Δ*lasR* co-cultures relative to each strain grown alone was not dependent on the initial ratio of WT to Δ*lasR* (**Fig. 1C**). We saw increased co-culture colony pigmentation when the initial proportion of the WT was at 0.2, 0.3, 0.5, 0.7, and 0.8 of the initial inoculum, with the balance comprised of Δ*lasR* (**Fig. 1C**). No increase in pyocyanin was observed at any ratio when WT was co-cultured with the Δ*lasR* complemented derivative (Δ*lasR* + *lasR*) indicating that the phenomenon was dependent on the *lasR* mutation (**Fig. 1C**). To assess the relative abundances of WT and Δ*lasR* in co-culture, we competed each strain against a neutrally-tagged WT strain (PA14 *att::lacZ*). We found that Δ*lasR* increased in proportion after 16 h growth in colony biofilms regardless of the starting proportion whereas the proportions of tagged and untagged WT remained stable (**Fig. 1D**). We have previously shown that Anr activity is higher in Δ*lasR* strains and contributes to Δ*lasR* competitive fitness against WT *P. aeruginosa* in colony biofilms (53, 54), but Anr was not required for co-culture pyocyanin production by Δ*lasR* (**Fig. S2**).

Pyocyanin is a product regulated by quorum sensing (QS) through the transcription factors LasR, RhlR, and PqsR (55-57), and because of the cell-density dependent quorum-sensing regulation, it was important to assess the population size in co-cultures relative to monocultures. Total CFUs did not increase in WT / Δ*lasR* mixed cultures relative to either strain alone (**Fig. S1D**). Instead we found that WT / Δ*lasR* co-cultures had fewer CFUs when compared to WT monocultures on LB (**Fig. S1D**). Taken together, these data suggested altered behavior, rather than cell number, contributed to the increased phenazine profile of LasR-strains.

### *P. aeruginosa* WT induces Δ*lasR* RhlR/I-dependent signaling independent of WT produced autoinducer

In the canonical QS pathway, LasR regulates both PqsR and RhlR, and mutants lacking either of these regulators in a WT background have impaired pyocyanin production (58, 59). Both *pqsR* and *rhlR* were required for the production of pyocyanin by Δ*lasR* in co-culture with WT (**Fig. 2A**). To determine if co-culture increased RhlR- or PqsR-dependent signaling in Δ*lasR* strains upon co-culture with the WT, we fused *lacZ* to the promoters of *rhlI* and *pqsA*, which provide readouts of the activity of each respective regulator (25). To examine the effects of WT on Δ*lasR* QS, we examined the interactions between WT and Δ*lasR* in single-cell-derived colonies. To do so, suspensions containing ∼50 cells of WT with ∼50 cells of either Δ*lasR* P*rhlI*–*lacZ* or Δ*lasR* P*pqsA* – *lacZ* were spread on LB agar with the colorimetric β-galactosidase substrate X-gal. Intercolony distances and β-galactosidase activity in Δ*lasR* strains were measured. We found that P*rhlI* – *lacZ* activity was inversely correlated with distance to a WT colony, with a significant increase in Δ*lasR* P*rhlI* – *lacZ* activity (**Fig. 2B**). Pearson correlation analyses showed that 54% of the variability in Δ*lasR* P*rhlI* – *lacZ* activity could be explained by changes in the distance to a WT colony (p-value ≤0.0001). The increased activity due to activation of P*rhlI* – *lacZ* in Δ*lasR* was not observed in Δ*lasR*Δ*rhlR*, and close proximity to another Δ*lasR* P*rhlI* – *lacZ* colony did not affect promoter activity (**Fig. 2B, inset**). Because RhlR activity is canonically stimulated by C4HSL (which is synthesized by RhlI) and P*rhlI* – *lacZ* is stimulated in Δ*lasR* by proximity to WT, we examined the role of RhlI in the Δ*lasR* response. We observed that a Δ*lasR*Δ*rhlI* strain was greatly impaired in the induction of pyocyanin upon co-culture with the WT (**Fig. 2A**) which suggests activation of RhlR involved C4HSL synthesis within Δ*lasR* strains. Although PqsR was required in Δ*lasR* for co-culture pyocyanin production, there was no significant correlation with proximity to WT for Δ*lasR* P*pqsA* – *lacZ* activity (**Fig. 2B**). Collectively, these data indicated that a diffusible factor produced by WT stimulated RhlR-dependent signaling in Δ*lasR* to induce downstream production of RhlR and PqsR dependent factors.

**Figure 2.**
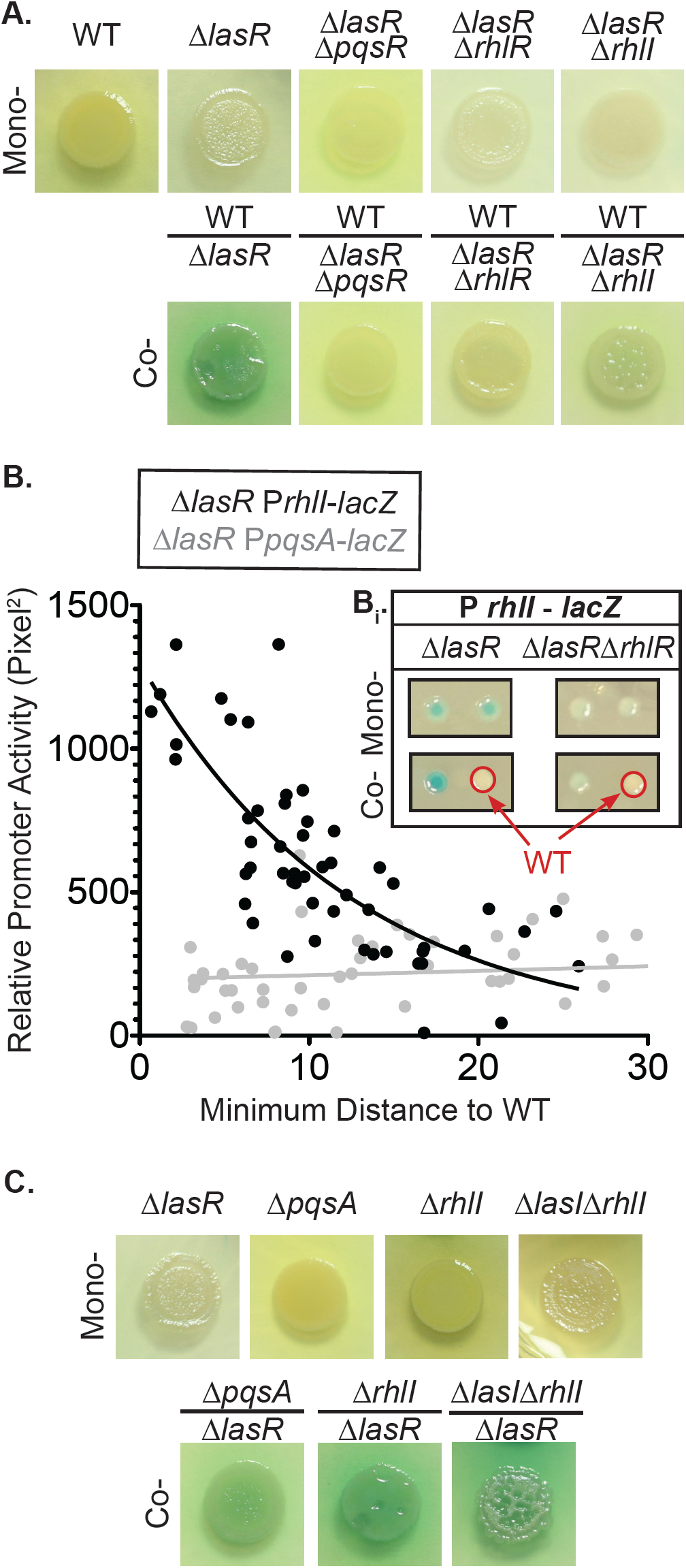
*P. aeruginosa* WT induces RhlR/I dependent activity in Δ*lasR* even in the absence of WT AHLs. **A**. Pyocyanin production by monocultures and WT co-cultures of Δ*lasR* and Δ*lasR* derivatives that are deficient PQS or RhlR/I quorum sensing on LB after 24 h growth. **B**. Promoter activity, quantified by relative pixel intensity of single cell-derived colony forming units (CFU) in co-culture with untagged WT CFU for Δ*lasR* P *pqsA* - *lacZ* (grey) and Δ*lasR* P *rhlI - lacZ* (black). Inset shows representative CFUs for RhlR-dependent Δ*lasR* P *rhlI* – *lacZ* activity when in monoculture and co-culture with WT (red circles). **C**. Co-culture pyocyanin production by Δ*lasR* still occurs upon co-culture with Δ*pqsA, ΔrhlI*, and *ΔlasI*Δ*rhlI* on LB after 24 h. All monocultures are shown at 48 h time point for comparison.

Because C4HSL is readily diffusible and likely produced by WT cells, we tested the hypothesis that C4HSL or other acylhomoserine lactones produced by WT were necessary to induce RhlR-dependent increases in *rhlI* promoter activity in Δ*lasR* co-cultured with WT. To test this hypothesis, we co-cultured Δ*lasR* with Δ*rhlI* or Δ*lasI*Δ*rhlI*, which lacks both acylhomoserine lactone synthases. Surprisingly, we found that like WT / Δ*lasR* co-cultures, Δ*rhlI / ΔlasR* co-cultures had higher levels of pyocyanin production relative to mono-cultures (**Fig. 2C**). Likewise, Δ*lasI*Δ*rhlI / ΔlasR* had higher levels of pyocyanin production relative to mono-cultures though the interaction was delayed by ∼24 h hours relative to the WT / Δ*lasR* co-cultures (**Fig. 2C**). Consistent with the data above which indicated that *pqsA* was not induced upon co-culture with the WT, Δ*lasR* co-cultures of the PQS-deficient strain Δ*pqsA* had high pyocyanin colony pigmentation relative to monoculture levels after 24 h of extended incubation (**Fig. 2C**). The dispensability of WT-produced quorum sensing signals implicated a novel signaling interaction in co-culture-dependent activation of Δ*lasR* RhlR-dependent signaling.

To assess whether RhlR activity in Δ*lasR* strains grown in the presence of WT was sufficient to elicit other RhlR/I-controlled phenotypes in addition to pyocyanin production, we tested whether co-culture with the WT also enhanced other RhlR-regulated processes in Δ*lasR*. RhlR regulates the production of rhamnolipid surfactants that are important for surface-associated motility known as swarming (60). While the rhamnolipid-defective mutant Δ*rhlA*, Δ*lasR*, and the Δ*lasR*Δ*rhlR* strains were not able to swarm, we observed increased swarming in co-cultures of Δ*lasR* with Δ*rhlA*. The phenomenon was dependent on RhlR as the Δ*lasR*Δ*rhlR* / Δ*rhlA* co-cultures did not swarm (**Fig. S3**). Altogether, these data suggested broad RhlR-mediated quorum sensing activation in LasR-strains grown in co-culture with LasR+ *P. aeruginosa.*

### Pyochelin production by Δ*lasR* is required for co-culture interactions

With evidence indicating that the induction of RhlR activity in Δ*lasR* in both mixed strain spot colonies and adjacent colonies by WT occurred through a mechanism that does not require acylhomoserine lactone cross-feeding, we sought to gain further insight into the mechanisms that underlie the co-culture interactions. We investigated the transcriptomes of the *lasR* mutant when co-cultured with either WT or itself via RNA sequencing analysis. We grew *ΔlasR* colony biofilms on LB physically separated from a lawn of either Δ*lasR* or WT by two 0.22 µm filters to prevent mixing of genotypes while allowing for the passage of small molecules. RNA was extracted from cells within the Δ*lasR* colony biofilms grown on the topmost filter for 16 h in order to examine Δ*lasR* transcriptional profiles (**Fig. 3A**). One hundred and ninety-nine genes in Δ*lasR* were higher and 198 genes were lower by a |log _2_ (fold change)| ≥ 1 with an FDR corrected p-value < 0.05 in co-culture with WT compared to Δ*lasR* alone (**Supplemental Table 1**). GO term analyses through PantherDB indicated that the upregulated gene set was significantly enriched in two pathways related to siderophore biosynthesis: pyoverdine biosynthetic process and salicylic acid biosynthetic process (an upstream precursor of pyochelin) with ∼44 and ∼77-fold enrichment, respectively (p-values < 0.005). Twenty-eight out of the 33 genes in the pyochelin and pyoverdine siderophore biosynthesis- and acquisition-related GO pathways were significantly upregulated in Δ*lasR* upon co-culture with WT (**Fig. 3B**). Other genes regulated by the low iron response were also differentially expressed including the *has* genes involved in heme uptake and *antABC* genes (**Table S1**). While we observed stimulation of *rhlI* promoter activity and increased production of RhlR regulated products, we did not see a pattern indicative of RhlR activation at this early time point (**Table S1**), and this point is discussed in more detail with respect to additional data shown below.

**Figure 3.**
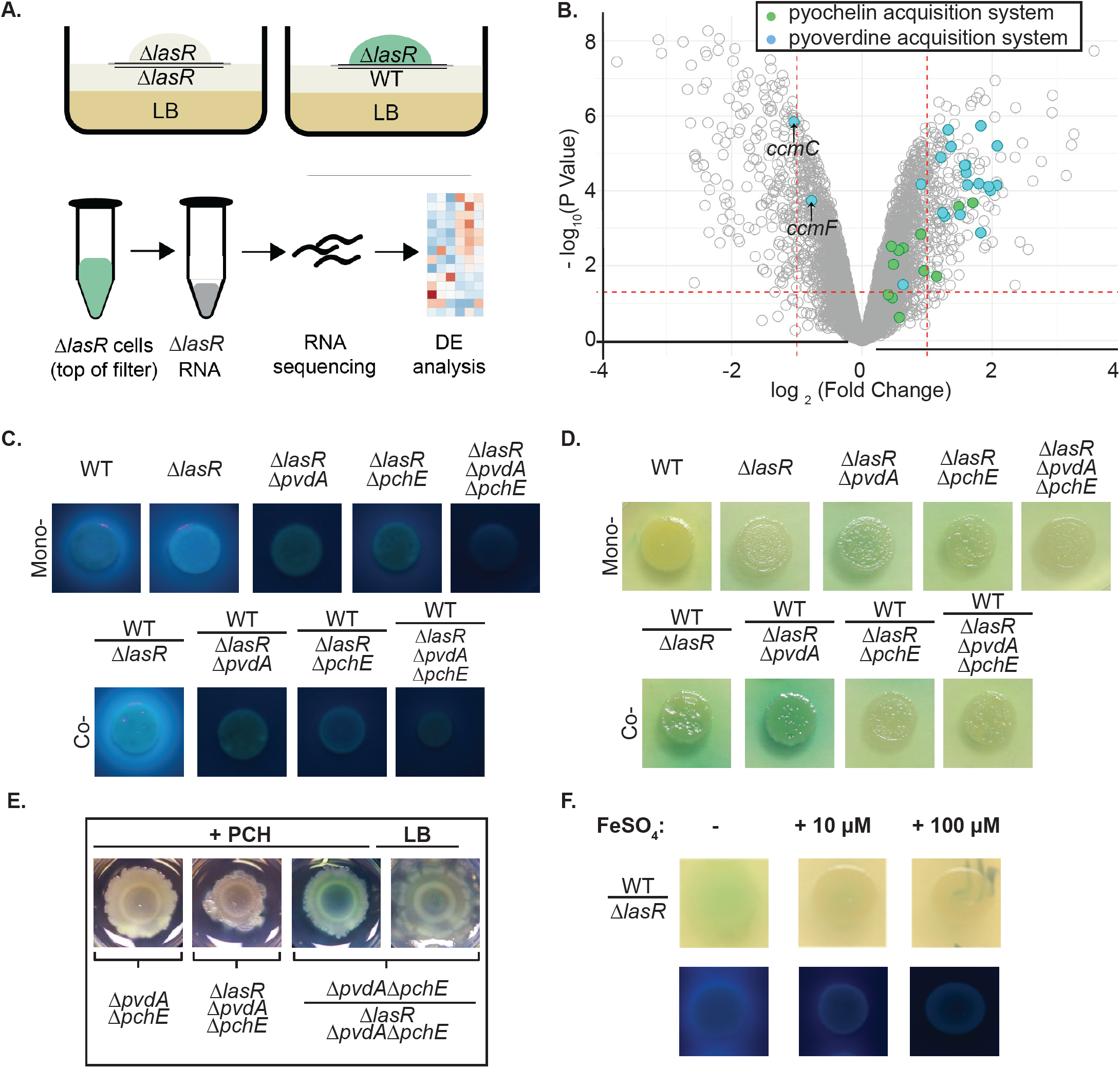
Biosynthesis of the co-culture-induced iron scavenging siderophore pyochelin is required in Δ*lasR* for pyocyanin over-production when cultured with wild type (WT). **A**. Scheme for the collection of RNA from Δ*lasR* colony biofilms grown above a lawn of WT or Δ*lasR*. **B**. Volcano plot of Δ*lasR* expression data with each point representing the log_2_(Δ*lasR* grown on WT / Δ*lasR* grown on Δ*lasR*) expression and – log_10_(P Value) of a single gene. Genes involved in pyoverdine (blue) and pyochelin (green) iron acquisition systems are indicated. *ccmC* and *ccmF* (indicated with arrows) of the pyoverdine GO term are involved in c-type cytochrome biosynthesis, and strains with knockouts of these genes are reported to produce more pyochelin. **C**. Mono- and co-cultures with Δ*lasR* strains deficient in pyoverdine (Δ*pvdA*) and/or pyochelin (Δ*pchE*). Colonies are visualized under ultraviolet light (UV) in order to see fluorescent siderophores. **D**. Pyocyanin production visualized for the colonies shown in panel **C. E**. Pyocyanin production by siderophore deficient strains grown in mono- and co-culture on LB with (+PCH) or without (LB) pyochelin-containing extract. Colonies were grown in a 12 well plate and imaged after 48 h. **F**. Wild-type and Δ*lasR* mixed colony biofilms grown on LB (-) or LB supplemented with either 10 or 100 µM FeSO4 visualized under ambient (top) and UV (bottom) light.

In light of the upregulation of siderophore biosynthesis genes in Δ*lasR* co-cultured with WT but not with itself, we qualitatively examined production of fluorescent pyoverdine and pyochelin siderophores in monocultures and co-cultures. To determine the contribution of both pyoverdine and pyochelin to fluorescence, genes required for pyoverdine biosynthesis (Δ*pvdA*), pyochelin biosynthesis (Δ*pchE*) or both pathways (Δ*pvdAΔpchE*) were disrupted in the *lasR* mutant (**Fig. 3C**). Increased fluorescence attributable to both pyoverdine and pyochelin in co-culture was due to siderophore production by Δ*lasR* strains, consistent with the RNA-Seq data, as the increased siderophore production in WT/ Δ*lasR* co-cultures was lost for co-cultures in which the Δ*lasR* strains were replaced with *ΔlasRΔpvdA*, Δ*lasRΔpchE*, or *ΔlasRΔpvdAΔpchE*. While WT and the pyoverdine-deficient derivative Δ*lasRΔpvdA* (i.e. WT / Δ*lasRΔpvdA*) showed increased pyocyanin production relative to either monoculture, Δ*lasRΔpchE* and *ΔlasRΔpvdAΔpchE* did not support the overproduction of pyocyanin in co-culture with WT (**Fig. 3D**). The decrease in Δ*lasR*-derived pyocyanin was not due to decreased fitness as disruption of *pvdA* and *pchE* in Δ*lasR* individually had no effect on the final proportions; in contrast, Δ*lasRΔpvdAΔpchE* had a significant defect in competitive fitness compared to the Δ*lasR* parental strain (**Fig. S4**). These data suggested that pyochelin played a role in the co-culture interaction. To test this model, we complemented the Δ*lasRΔpchE* in co-culture with supernatant extracts from cultures of Δ*pvdA* which only produced pyochelin, or Δ*pvdAΔpchE* which produced neither siderophore (**Fig. S5A** for supernatant absorption spectra). The two supernatants were analyzed using the chrome azurol S (CAS) assay (61) to confirm that chelator activity was present in the Δ*pvdA* supernatant extracts but not in extracts from Δ*pvdAΔpchE* cultures (**Fig. S5B**). The amendment of the medium with PCH-containing extracts, but not siderophore-free extracts, restored co-culture pyocyanin production in Δ*pvdAΔpchE* / Δ*lasRΔpchE* co-cultures (**Fig. 3E**) lending further support for the model that pyochelin was required for co-culture interactions. Consistent with the requirement of the siderophore for the stimulation of pyocyanin in WT / Δ*lasR* co-cultures, iron supplementation of the LB medium suppressed siderophore production and the stimulation of pyocyanin production (**Fig. 3F**).

### Evidence for WT - Δ*lasR* interactions involving citrate secreted by the WT in response to pyochelin

Many of the genes with higher transcript levels in Δ*lasR* upon co-culture with WT have annotations related to organic acids such as anthranilate and citrate (**Supplemental Table 1, Fig. S6A**). First, anthranilate was examined as a candidate molecule that was produced by the WT and that induced RhlR-dependent phenotypes in Δ*lasR*. Co-cultures of Δ*lasR* with anthranilate synthase mutant Δ*phnAB* did not alter high phenazine production compared to WT / Δ*lasR* (**Fig. S6B**). Additionally, anthranilate supplementation of the medium did not alter Δ*lasR* phenazine production at any concentration tested up to ∼15 mM (**Fig. S6C**). Furthermore, co-cultures of Δ*lasR* and the PQS deficient strain Δ*pqsA*, which accumulates anthranilate as an upstream precursor (62), did not show enhanced stimulation of RhlR-dependent phenotypes beyond WT / Δ*lasR* levels (**Fig. 2C**).

We then tested the ability of citrate to induce RhlR-dependent phenotypes in Δ*lasR* in light of the observation that twenty percent of the most strongly differentially expressed genes (|log _2_ (fold change)| ≥ 2 with an FDR-corrected p-value < 0.05) were implicated with citrate sensing, transport, catabolism, and anabolism as annotated by UNIPROT and pseudomonas.com (**Fig. 4A**). Among the genes more highly induced in Δ*lasR* / WT co-cultures compared to Δ*lasR* cultured with itself were genes annotated as playing a role in citrate sensing or metabolism, with the most strongly upregulated genes were involved in citrate catabolism and sensing and transport (**Fig. 4B**).

**Figure 4.**
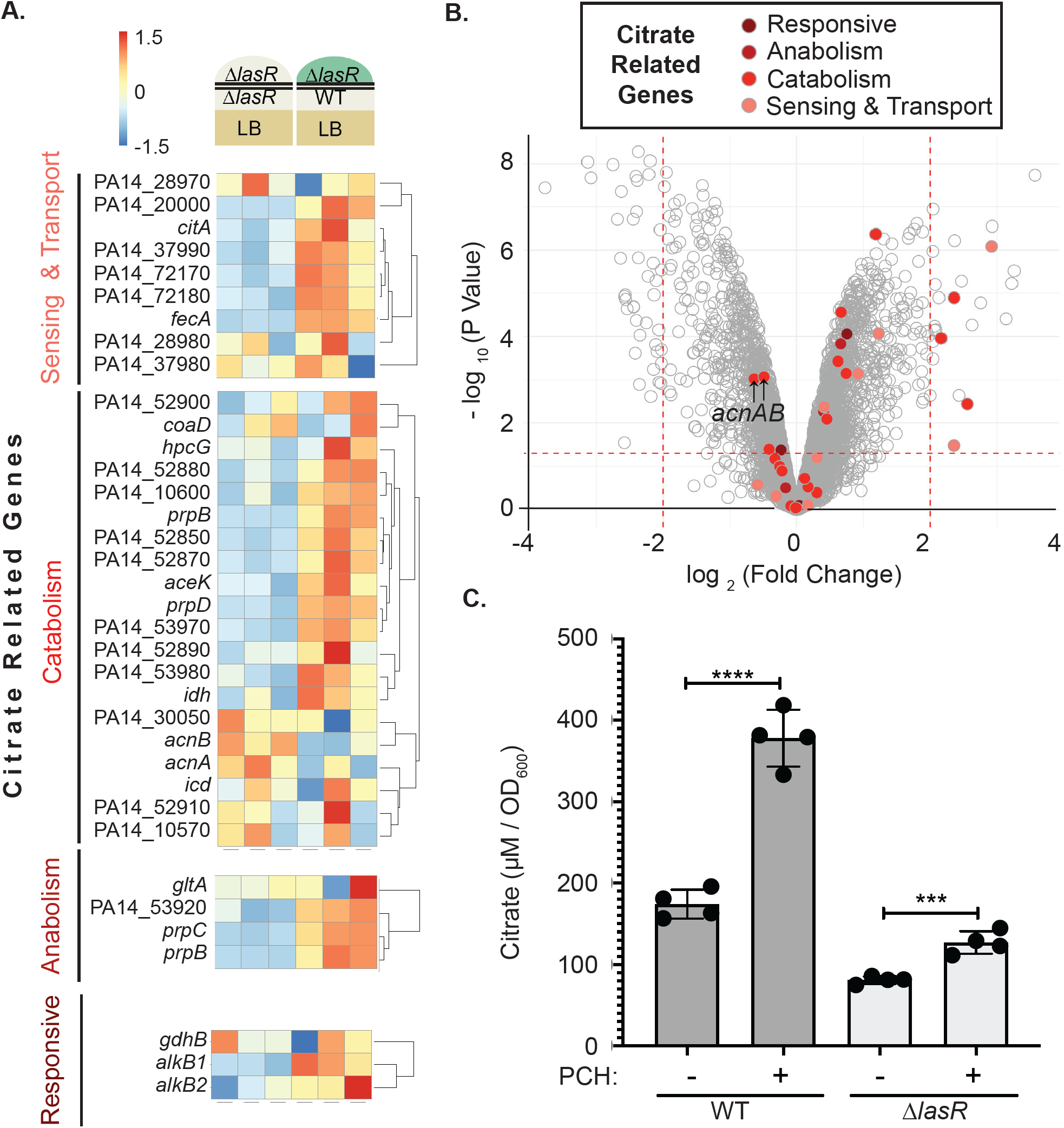
Citrate release by WT is induced by pyochelin exposure. **A**. Subset of Δ*lasR* co-culture expression data (see **Fig. 3A** for set up) for genes annotated as being involved in citrate sensing, transport, catabolism, anabolism, and those shown to be responsive to citrate. **B**. Volcano plot of Δ*lasR* co-culture expression data with genes shown in panel A highlighted in red. **C**. Citrate concentrations in supernatants from wild type and Δ*lasR* stationary phase cultures after growth in LB supplemented with extracts containing 50 µM pyochelin (PCH +) or an equal volume of control extracts (PCH -). A representative experiment with four biological replicates is shown; ***, p≤ 0.001 and ****, p≤ 0.0001 by two-tailed t-test of paired ratios.

In light of the findings that Δ*lasR* strains induced low iron responsive genes when grown near the WT but not itself, that Δ*lasR* pyochelin production was necessary for co-culture interactions that lead to increased pyocyanin and *rhlI* promoter activity, that citrate sensing and catabolism genes were induced in Δ*lasR* by the presence of the WT, and the knowledge that numerous microbes, including *Pseudomonas putida*, secrete citrate and other organic acids when iron limited (49, 50, 63, 64), we measured citrate in the supernatants of WT and Δ*lasR* LB cultures. Citrate concentrations were significantly higher in WT supernatants (**Fig. 4C**). Furthermore, amendment of the LB medium with extracts containing 50 µM pyochelin increased extracellular citrate concentrations by about 2-fold in WT cultures compared to control cultures containing extracts lacking pyoverdine and pyochelin, with a much smaller stimulation in Δ*lasR* cultures (**Fig. 4C**). The difference in the effects of pyochelin on citrate levels in culture supernatants for WT and Δ*lasR* was significant using data from four independent experiments (p<0.05). This suggested WT-produced citrate was possibly involved in WT / Δ*lasR* co-culture interactions, and that its increased release was enhanced by Δ*lasR*-produced pyochelin.

### Citrate induces RhlR activity in Δ*lasR* and RhlI levels in a ClpX-protease dependent manner

To determine if citrate was sufficient to stimulate RhlR activity in Δ*lasR*, we analyzed its effects on both *rhlI* promoter fusion activity and RhlI protein levels. We found that citrate increased *rhlI* promoter activity in Δ*lasR* and that its effects were dependent on the presence of RhlR (**Fig. 5A**). In contrast, the inclusion of citrate in the medium caused a small but significant reduction in WT P*rhlI* activity compared to LB control (**Fig. 5A**).

**Figure 5.**
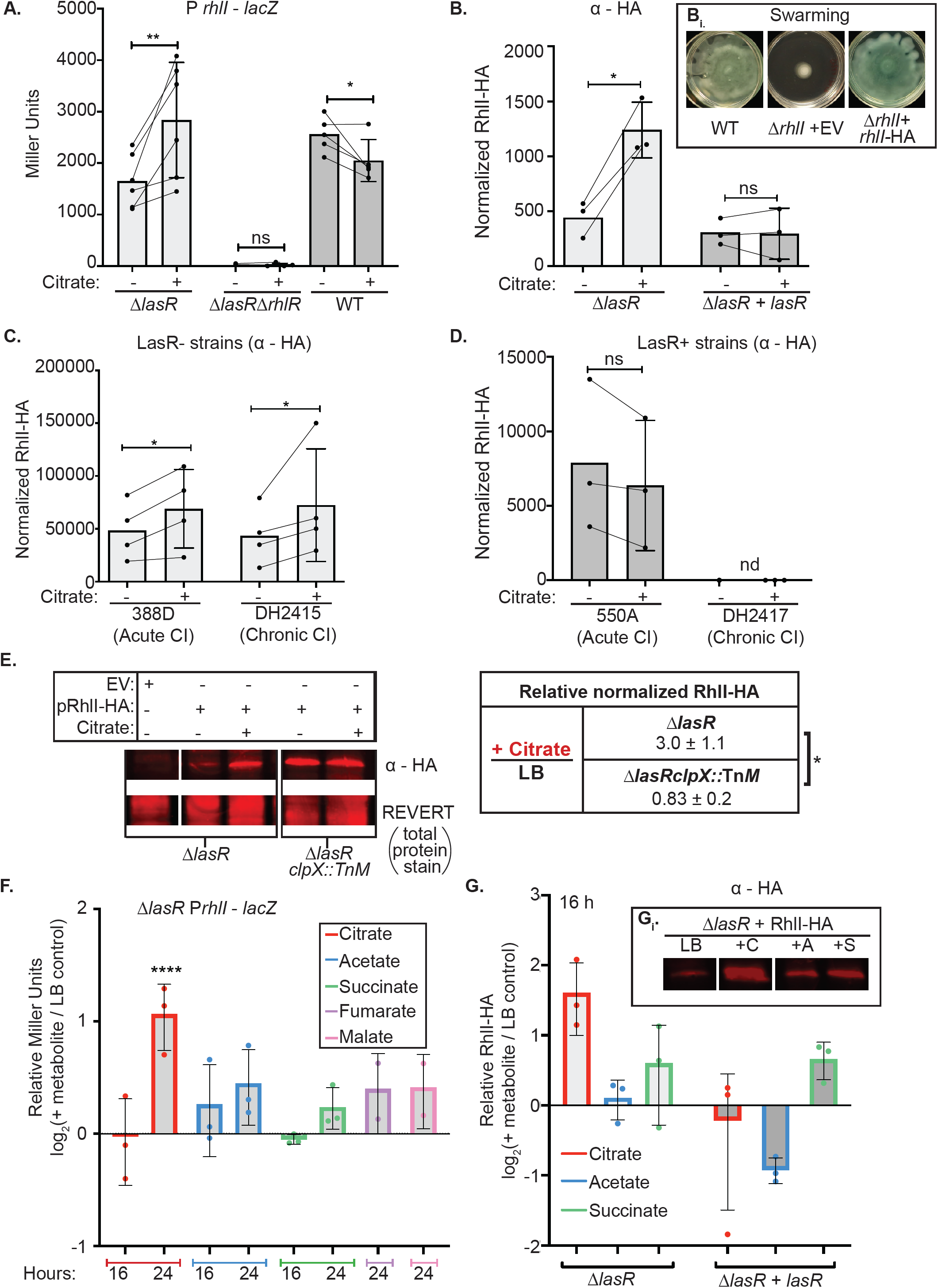
Citrate and related compounds induce RhlR-dependent *rhlI* promoter activity and stabilize RhlI protein in LasR-strains in a ClpX protease dependent manner. **A**. β-galactosidase activity for Δ*lasR, ΔlasRΔrhlR*, and wild type harboring *att::*P*rhlI - lacZ* on LB ± 20 mM citrate at 24 h. Each point is the average of three biological replicates from 3 - 4 independent experiments. Statistical analyses performed by one-way ANOVA with Dunnet’s multiple hypotheses correction. **B**. RhlI-HA protein signal normalized to REVERT total protein stain (Licor) on LB ± 20 mM citrate for Δ*lasR* and *lasR* complemented at the native locus (Δ*lasR*+*lasR*). n = 3 biological replicates performed on three independent days. Dunnet’s multiple comparison test with LB control. Inset illustrates that plasmid-borne RhlI-HA, but not the empty vector (EV) can complement an Δ*rhlI* mutant for swarming. **C**. RhlI-HA protein levels on LB and LB supplemented with 20 mM citrate of LasR LOF (LasR-) clinical isolates (CI) from acute corneal (388D) and chronic CF infections (DH2415). **D**. RhlI-HA protein levels on LB ± 20 mM citrate of LasR+ acute corneal CI (550A) of same MLST type as 388D and LasR+ chronic CF CI (DH2417) from which DH2415 evolved. **E**. Representative image and quantification of replicates for the anti-HA antibody analysis of Δ*lasR* and Δ*lasRclpX*Tn::*M* + pRhlI-HA or pEV grown in LB and LB supplemented with 20 mM citrate. **F.** RhlR-dependent *rhlI* promoter activity of Δ*lasR* P *rhlI - lacZ* on LB supplemented with citrate, acetate, succinate, fumarate, and malate at 16 and 24 h relative to LB control. **G**. Normalized RhlI-HA protein levels Δ*lasR* and Δ*lasR* + *lasR* at 16 h of growth on citrate, acetate, and succinate. Inset shows representative blot of RhlI-HA protein under these same conditions. p-values: * (p<0.05), ** (p ≤ 0.05), *** (p≤ 0.001), and **** (p≤ 0.0001).

To determine if RhlI protein levels were influenced by citrate, we utilized an arabinose-inducible *rhlI-*HA construct to assess RhlI protein levels and stability of RhlI-HA in the absence and presence of citrate. RhlI-HA was functional as swarming defects of Δ*rhlI* were complemented upon expression of RhlI-HA but not by the empty vector (**Fig. 5B, inset**). RhlI-HA protein levels were 3-fold higher in Δ*lasR* upon citrate supplementation relative to controls (**Fig. 5B**). Consistent with the absence of an increase in *rhlI* promoter activity in WT strains (**Fig. 5A**), RhlI-HA protein levels were not higher with citrate in the Δ*lasR* complemented strain (Δ*lasR* + *lasR*) (**Fig. 5B**). The differential responses to citrate were also observed in LasR- and LasR+ pairs of clinical isolates (CIs). LasR-CIs from acute (strain 388D) and chronic (strains DH2415) infections had RhlI-HA levels 1.4- and 1.7-fold higher, respectively, in the presence of citrate (**Fig. 5C**), whereas alterations in RhlI-HA protein levels in LasR+ CIs from acute (550A) or chronic (DH2417) infections was not observed (**Fig. 5D**). Through this work, we successfully identified citrate as a molecule in co-culture that specifically promoted RhlI protein levels in LasR-strains, but not LasR+ strains, by a mechanism other than transcriptional control. In order to identify transporters that could be involved in the Δ*lasR* response to citrate, we deleted two organic acid transporters: *dctA* (65) and PA14_51300 (66) in the Δ*lasR* background. The *dctA* gene was deleted in both a Δ*lasR* and Δ*lasR*Δ*rhlR* mutant. We found that the Δ*lasR*Δ*dctA* strain still showed induction of pyocyanin when co-cultured with the WT and that induction was dependent on RhlR (**Fig S7A**). Similar results were obtained with the Δ*lasR*ΔPA14_51300 (**Fig S7B**) suggesting that these transporters were not required for the interaction perhaps due to redundant functions of other proteins or the involvement of other import mechanisms.

The temporal pattern suggested RhlI protein induction preceded signal amplification via the positive feedback loop of the quorum sensing transcriptional network. This would be consistent with a primary effect on post transcriptional modulation of RhlI-mediated RhlR activity. To begin to unravel the mechanisms by which citrate promoted RhlR/I-dependent signaling and RhlI stability in Δ*lasR*, we analyzed the role of two proteases previously found to target and degrade RhlI (i.e. Lon and ClpXP) (67). Given knockouts of Lon protease have a less substantial rise in RhlR/I expression in Δ*lasR* knockouts compared to WT (68), we focused on the role of ClpXP in Δ*lasR*. We found that citrate induction of RhlI-HA protein levels in Δ*lasR* relative to the LB control was dependent on the production of ClpX protease (**Fig. 5E**). More specifically, in the absence of ClpX, a protease shown to degrade RhlI (i.e. Δ*lasRclpX*::Tn*M*), RhlI-HA levels did not increase on LB + citrate relative to LB control, unlike Δ*lasR* comparator (**Fig. 5E**). In LB conditions, RhlI-HA levels were 3.20 ±2.1 fold higher in Δ*lasRclpX*::Tn*M* compared to Δ*lasR*, which mirrors the 3-fold induction observed for Δ*lasR* on LB + citrate. No significant difference in RhlI-HA were observed for Δ*lasRclpX*::Tn*M* relative to Δ*lasR* in citrate supplemented conditions (fold change: 1.01 ±0.53). In other words, as previously noted for WT strains, ClpX appeared to degrade RhlI in Δ*lasR*, and played a role in Δ*lasR* response to citrate. The distinct responses and mechanisms identified between LasR+ and LasR-strains under iron limitation and exposure to the low-iron associated molecules, citrate and pyochelin, enabled increases in antagonistic factor production beyond monoculture levels as an emergent property of *P. aeruginosa* intraspecies interactions.

In support of the model that induction in RhlR signaling in response to citrate was due to increased RhlI protein, we found that the induction of *rhlI* promoter activity (**Fig. 5F**) followed the increase in RhlI-HA levels (**Fig. 5G**) in response to citrate. The stimulation of *rhlI* promoter activity was greatest for citrate, but modest stimulation was observed for other organic acids including TCA cycle intermediates (succinate, fumarate and malate) and the common fermentation product acetate. In each case, the stimulation of *rhlI*-promoter activity was accompanied by higher RhlI-HA protein levels in Δ*lasR*, but not Δ*lasR + lasR*, aside from succinate (**Fig. 5F**,**G**). Together, these data may imply that organic acids, such as citrate, can serve as mediators of co-culture interactions that can activate RhlR activity in LasR-strains.

## Discussion

In this study, we described an emergent outcome of co-culturing LasR- and LasR+ strains of *P. aeruginosa* in which their interactions promoted the increased production of toxic exoproducts including pyocyanin and rhamnolipids (see **Fig. 6** for model). We determined that, in co-culture, the iron-binding siderophore pyochelin was largely contributed by Δ*lasR*, and that exogenous pyochelin induced secretion of citrate significantly more strongly in the WT than in Δ*lasR*. Citrate increased RhlI protein levels and activated RhlR activity only in Δ*lasR*, but not WT cells (**Fig. 6**). Western blot analysis of RhlI-HA expressed from a regulated promoter led us to propose that the increase in RhlR signaling is due to decreased degradation of RhlI by ClpXP, a known negative regulator of RhlI (68, 69). The differences in siderophore production, citrate release, and RhlR/I activation between *P. aeruginosa* LasR+ and LasR-strains in co-culture environments reflect the pronounced differences between strains that drive the reactivation of quorum sensing and enhanced production of secreted factors. Previous studies have shown that LasR-strains increase their production of phenazines in the presence of other species such as *Candida albicans* (22) and *Staphylococcus aureus* (see Fig. 3B in (70)) and future work will determine if pyochelin and citrate also participate in these interspecies interactions. Other microbial interactions have been shown to be influenced by iron availability (71-74). Furthermore, the activation of RhlR activity in Δ*lasR* strains that can occur in late stationary phase cultures (23, 75) may relate to changes in iron or TCA cycle intermediates. While we found that WT production of the diffusible autoinducers 3OC12HSL, C4HSL and PQS were not required for co-culture stimulation, they clearly contributed to the enhanced activation of RhlR regulation which is consistent with intercolony QS interactions that have been demonstrated previously (76).

**Figure 6.**
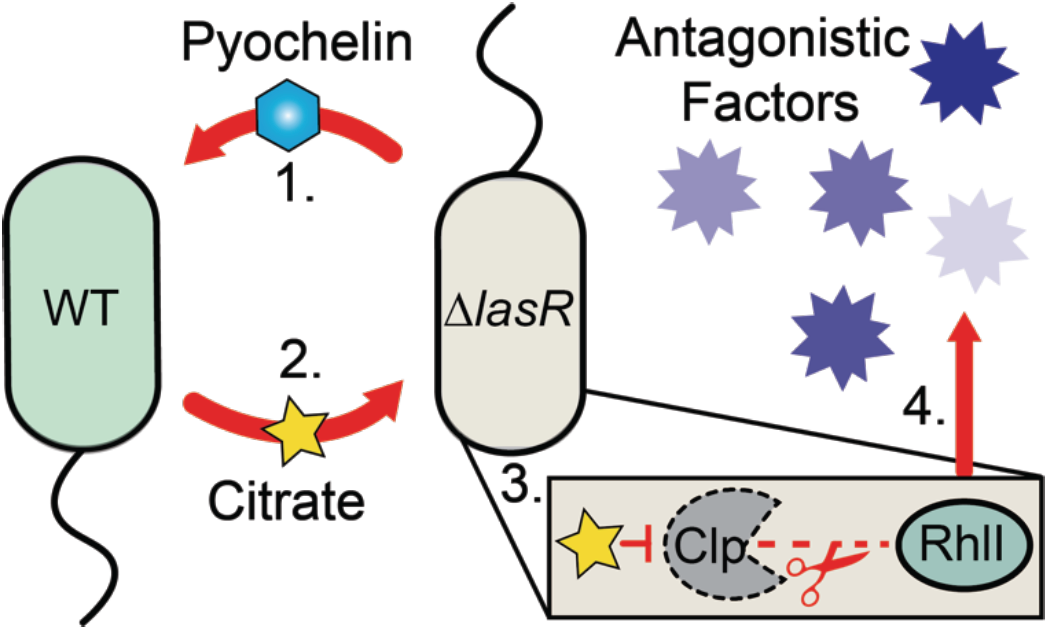
Model for wild type and Δ*lasR* co-culture interactions. **(1.)** Δ*lasR* produced pyochelin promotes citrate release in wild type. **(2.)** Citrate released by wild type in co-culture stimulates RhlR/I dependent activity **(3.)** by stabilizing RhlI protein in ClpXP protease dependent mechanism **(4.)** to promote the production of antagonistic factors like pyocyanin toxin and rhamnolipid surfactant.

The stimulatory relationship between LasR+ and LasR-strains was remarkably stable as it was observed when strains were mixed within single spot colonies (**Fig. 1A**) and when strains separated by either filters (**Fig. 3**) or mm distances on an agar plate (**Fig. 2B**). The LasR-/ LasR+ interactions occurred across distinct media (**Fig. S1A**), among genetically diverse LasR+ and LasR-clinical isolates (**Fig. S1B**) and over a wide range of relative proportions of each type (**Fig. 1C**). The consequences of this intraspecies interaction between genotypes may explain the worse outcomes exhibited by patients in which LasR-strains are detected (12), but future studies with data that include genotypes, mono-culture and co-culture phenotypes, and longitudinal outcome data will be required. RhlR plays other important roles in host interactions (77) which may benefit *P. aeruginosa* LasR-strains. The observation that *rhlR* mutants are rare in natural *P. aeruginosa* isolates and that LasR-strains with active RhlR are virulent (21, 78) underscores the relevance of this mechanism and highlights the importance of understanding how microbial interactions activate RhlR.

As the study of inter- and intra-species interactions progresses, it is becoming increasingly clear that the environment can dictate the outcome of microbial interactions (79). In fact, even the importance of QS regulation for fitness depends on nutrient sources and conditions (80, 81). As Δ*lasR*-produced pyochelin was a key component of the interaction, and pyochelin production is repressed under conditions of excess iron availability, it was not surprising that the addition of iron to LB medium suppressed the interaction without significantly altering the final colony CFUs or strain ratios relative to LB control (**Fig. S4**). Siderophore-mediated iron uptake is often required *in vivo* (39, 82, 83) due to iron sequestration by host proteins (84-87), thus the *in vivo* settings could support these interactions. Interestingly pyoverdine, the higher affinity siderophore, was not required for the co-culture response mirroring findings that genes for the biosynthesis of pyoverdine, but not pyochelin, are commonly disrupted in chronic CF clinical isolates (44-46). In the absence of pyoverdine (i.e. Δ*lasRΔpvdA*), we observed more pyocyanin in co-culture with WT than Δ*lasR* (**Fig. 3D**), and we speculate that this is due to increased pyochelin production by Δ*pvdA* but future studies will be required to test this model. If this is the case, it would be interesting to analyze the outcomes of interactions over gradients of iron and other nutrients. It was interesting to find that in WT / Δ*lasR* co-culture, heme-related proteins, *hasAP, hasS*, and *hasD*, were among the top eight most upregulated genes by Δ*lasR* because the presence of *lasR* mutants and heme utilization are both reported biomarkers of disease progression in CF patients (12, 88). Co-culture induced *lasR* mutant phenotypes may link these two correlative observations.

Citrate, a TCA intermediate, is released under iron limitation as a result of “overflow metabolism” (48-50, 52) and is also used by *P. aeruginosa* and other microbes for iron acquisition due to its iron chelating properties (89). The higher siderophore production by Δ*lasR* and stimulation of Δ*lasR* siderophore production in co-culture likely reflects different metabolic strategies between the two strains. Ongoing work will investigate the mechanisms that drive differences in metabolism and iron requirements in order to determine how these differences shape microbial and host interactions. It is likely that Crc-mediated catabolite repression is involved in the response to citrate and the control of RhlI levels (67, 69). The existence of a mechanism for the induction of RhlR-mediated QS in response to citrate and other TCA cycle intermediates that are secreted when iron is limiting dovetails with reports of increased expression of the *P. aeruginosa* quorum sensing regulon in low iron in LasR+ cells (90-92). This coordinate regulation may aid in iron acquisition as quorum sensing-controlled phenazines, such as pyocyanin, reduce poorly soluble Fe_3+_ to Fe_2+_ and facilitate its uptake via the Feo system (93). Additionally, rhamnolipids have been employed for iron remediation (94, 95) which suggests their surfactant activity may increase *P. aeruginosa* substrate iron uptake in part through hydroxyalkylquinolone-dependent mechanisms (96).

As the presence of heterogeneous genotypes within single species populations becomes increasingly appreciated, it is important to understand how commonly encountered genotypes interact to influence the apparent behavior of the population. Here, we show that inter-genotype interactions lead to increased RhlR signaling in *lasR* strains; other work shows co-cultures can also influence the survival of other genotypes (97). It is likely that a wide array of such interactions have yet to be uncovered.

## Methods

### Strains and Growth Conditions

Bacterial strains used in this study are listed in **Table S2**. Bacteria were maintained on LB (lysogeny broth) with 1.5% agar. Yeast strains for cloning were maintained on YPD (yeast peptone dextrose) with 2% agar. Where stated, 20 mM of indicated metabolite was added to the medium (liquid or molten agar). Planktonic cultures were grown on roller drums at 37°C for *P. aeruginosa*.

### Plasmid Construction

Plasmid constructs for making in-frame deletions, RhlI-HA expression, and for *pqsA* promoter fusions were constructed using a *Saccharomyces cerevisiae* recombination technique described previously (98). The RhlI-HA expression vector with the ampicillin cassette (pMQ70) was constructed by amplifying the *rhlI* gene with primers that added an HA-tag with sites for cloning into pMQ70 (AmpR). All plasmids were sequenced at the Molecular Biology Core at the Geisel School of Medicine at Dartmouth. In frame-deletions and integrated promoter fusions were introduced into *P. aeruginosa* by conjugation via S17/lambda pir *E. coli*. Merodiploids were selected by drug resistance and double recombinants were obtained using sucrose counter-selection and genotype screening by PCR. Both RhlI-HA expression vectors were introduced into *P. aeruginosa* by electroporation. The Δ*lasRclpX*::Tn*M* was identified from a collection of transposon mutants in the Δ*lasR* and verified by sequencing.

### Pyocyanin Quantification

*P. aeruginosa* strains were grown in a 96-well plate containing 200 µL LB agar per well by inoculation with 5 µL of overnight cultures adjusted to OD_600_ = 1. After 16 h incubation at 37 °C, two agar plugs (with indicated *P. aeruginosa* mono- or co-cultures) were added to tubes containing chloroform (500 µL), mixed by vortexing for 30 s and then centrifuged for 2 min at 13,000 RPM. The lower chloroform layer (200 µL) was collected into new tubes, and the chloroform extraction was repeated with an additional 500 µL of chloroform. The chloroform extracts (400 µL) were acidified with 0.2 N HCl (500 µL) and vortexed for 30 s. The pink aqueous layer containing pyocyanin was diluted 1:2 in 200 mM Tris-HCl (pH 8.0). Relative pyocyanin was measured by reading absorbance at 310 nm relative to media blank extracts. Values were reported per plug. Each condition had at least eight replicates each for two independent experiments.

### Competition Assays

Competition assays were performed to determine the relative fitness of *P. aeruginosa* mutants. Strains to be competed were grown overnight and adjusted to OD_600_ = 1. Competing strains were combined with PA14*att*::*lacZ* strain in a 1:1 ratio, unless otherwise stated. Following 15 s vortex, 5 µL of the combined suspension was spotted on LB agar. After 16 h, colony biofilms (and agar) were cored, placed in 1.5 mL tubes with 500 µL LB, and agitated vigorously for 5 min using a Genie Disruptor (Zymo). This suspension was diluted, spread on LB plates supplemented with 150 µg / mL 5-bromo-4-chloro-3-indolyl-D-galactopyranoside (X-Gal) using glass beads, and incubated at 37 °C until blue colonies were visible (∼24 h). The number of blue and white colonies per plate were counted and the final proportions recorded. Each competition was run in triplicate on 3 separate days.

### Colony Proximity Image analysis

Glass beads were used to spread 50 µL of a 1:1 mixture of untagged WT and Δ*lasR* possessing the indicated promoter fusion to *lacZ* onto LB plates (2% agar) supplemented with 150 µg / mL 5-bromo-4-chloro-3-indolyl-D-galactopyranoside (X-Gal). After 16 h incubation at 37 °C, plates were placed at 4 °C for an additional 24 h to allow all Δ*lasR* colonies to develop blue coloration of various intensities. Plates were imaged on glass sheet to reduce glare using Canon EOS Rebel T6i digital camera. To process images, they were first cropped to remove background area surrounding each plate, converted to 8-bit for particle analysis in ImageJ, and the threshold was determined to count WT and Δ*lasR* CFU’s, separately. Colony parameters were collected for each individual CFU, including x, y coordinates and area for WT and Δ*lasR* CFU lists. A simple distance calculation was made between every WT and Δ*lasR* colony using the x,y coordinates and the minimum distance to a WT colony was plotted for each Δ*lasR* CFU against its area value, representative of the approximate *lacZ* intensity.

### Swarming Motility Assays

Swarm assays were performed as previously described in (99), with a few modifications. Briefly, M8 medium with 0.5 % agar was poured into 60 x 15 mm plates and allowed to dry at room temperature for 4 h prior to inoculation. LB grown cultures (16 h at 37 °C) were diluted to OD_600_ = 1 in fresh LB, and co-cultures were mixed such that Δ*rhlA* was at 0.7 proportion of the final cell suspension. Each plate was inoculated with 5 µL of the final cell suspensions and incubated upright for 24 h at 37 °C in an incubated chamber followed by 12 – 16 h at room temperature. Each strain was inoculated in four replicates and assessed on at least three separate days.

### RNA Collection

A 200 µL aliquot of optical density normalized (OD_600_ = 1) cultures of PA14 or PA14 Δ*lasR* from three independent overnights were spread onto LB plates with glass beads and briefly allowed to dry. Two isopore 0.2 µm PC membrane filters were stacked on the lawn (37 mm diameter filter directly on lawn then 25 mm diameter filter on top), and three 15 µL spots of normalized (OD600 = 1) Δ*lasR* cultures were spotted on the top-most filter. After 16 h incubation at 37 °C, the top filter was collected and Δ*lasR* cells were resuspended in 1 mL LB by 5 min of vigorous shaking on the genie disruptor. Cells were pelleted for 10 min at 13,000 RPM and snap-frozen in ethanol and dry ice for RNA extraction. RNA was extracted according to manufacturer’s protocol with the QIAGEN RNeasy Mini kit and DNase treated twice with Invitrogen Turbo DNA-Free kit. DNase-treated samples were prepared for sequencing with ribodepletion and library preparation in accordance with Illumina protocols. Samples were barcoded and multiplexed in a NextSeq run by the Dartmouth Sequencing Core.

### RNA-Seq Processing

Reads were processed using CLC Genomics Workbench wherein reads were trimmed and filtered for quality using default parameters. Reads were aligned to the *P. aeruginosa* UCPBB_PA14 genome from www.pseudomonas.com. Results were exported from CLC including total counts, CPM and TPM. EdgeR was used to process differential gene expression (100). Generalized linear models with mixed effect data design matrices were used to calculate fold-change, p-value and FDR. Volcano plots and heatmaps using EdgeR output (log fold-change and -log(p-value)) were made in R (ggplot2 and pheatmap respectively) (101-103). GO term pathway enrichment analysis was carried out using PantherDB (104).

### Accession Number

Data for our RNA-Seq analysis of *P. aeruginosa* Δ*lasR* grown on Δ*lasR* or WT in co-culture has been uploaded to the GEO repository (https://www.ncbi.nlm.nih.gov/geo/) with the accession number GSE149385.

### Pyochelin extraction, quantification, and validation

Pyochelin was extracted based on the methods of Cox et al. (37). Briefly 50 mL cultures of PA14 Δ*pvdA* and pyochelin biosynthesis deficient strain PA14 Δ*pvdAΔpchE* (negative control) were grown in Chelex-treated (i.e. media treated for > 2 h with 0.5 g Chelex resin per 10 mL media concentrate, followed by centrifugation and filtration with 0.22 µm filter unit) LB for 16 h. Cultures were pelleted for 15 min at > 5000 RPM, and the supernatant was passed through a 0.22 µm filter unit. The cell-free supernatant was acidified to pH 2 with 10 N HCl. For extraction, 5 mL ethyl acetate was added per 50 mL acidified solution in a separatory funnel. The top ethyl acetate layer was concentrated using a speedvac and quantified in 50 / 50 methanol:dH2O in a 1 mm quartz cuvette at 313 nm. Absorbance was checked from 200 - 600 nm for expected peak profile. For some extractions, a 5 µL aliquot of Δ*pvdA* extract in 50:50 methanol:dH2O was viewed under ultraviolet light for expected fluorescence relative to Δ*pvdAΔpchE* extract that dissipated upon 10 µM FeSO4 supplementation. The concentration was determined using the molar extinction coefficient at 313 nm in 50 / 50 methanol:dH2O in a 1 mm quartz cuvette (37). Upon quantification, concentrated extract was lyophilized using rotovap to yellow resin, and used within 2 days of initial extraction by resuspension in LB for supplementation. As validation of biological activity, 10 µL of extract was spotted along with EDTA chelator as control on CAS agar prepared as described in.

### Citrate Quantification

Citrate was quantified from cell-free supernatant of 5 mL LB-grown cultures inoculated from single colonies. Cultures were grown in quadruplicate and incubated on a roller drum for 16 h. OD _600 nm_ was recorded, and cultures were pelleted for 15 min at > 5,000 RPM. The supernatant was passed through a 0.2 µm syringe filter unit. Citrate in the filtered supernatant was quantified according to manufacturer’s “manual assay” protocol (Megazyme) in ½ reactions. The extinction coefficient at 340 nm in a quartz cuvette was used to quantify concentration of citrate relative to OD _600_ at the end of the 5 min enzymatic reaction. Citrate standard and blank media conditions were included in every assay. The average for 4 replicates for each experiment was reported across 3 - 4 independent days.

### Beta galactosidase assays

Cells with a promoter fusion to *lacZ - GFP* integrated at the *att* locus were grown in 5 mL cultures of LB at 37°C for 16 h. The cultures were diluted to a starting OD _600_ of 1 and 5 µL were spotted onto LB agar plates ±20 mM pH 7 citrate (or other specified metabolite) in triplicate. After 24 h (or other indicated time) at 37 °C, colony biofilms were cored, resuspended in 500 µL LB by vigorous shaking on the Genie Disrupter for 5 min as previously described, and β-Gal activity was measured as described by Miller (105). The average for each experiment was reported across 3-4 independent days.

### Western Blot

Strains were grown in LB broth under selection (60 μg / mL gentamycin or 60 μg / mL carbenicillin as appropriate) for 16 h at 37 °C on a roller drum, and 5 μL of culture was spotted onto LB plates under selection with 0.2% L-arabinose (v/v) and +/- 20 mM indicated carbon source. Inoculated plates were incubated at 37 °C for 16 h. Colony biofilms were resuspended in 325 μL of Laemmli buffer without reducing agent and heated at 100 °C for 15 min. Protein was quantified on 1:10 dilution of protein sample according to standard procedure via Thermo scientific BCA Protein Assay Kit. Reducing agent was added and samples were run on a 4 - 15% SDS gradient gel (Bio-Rad) at 60 V for 40 min followed by 110 V for 45 min. After SDS page electrophoresis, protein was transferred onto LF-PVDF (Bio-Rad) using the mixed molecular weight option on a turbo blot apparatus (Bio-Rad). After transfer the membrane was dried, rehydrated, and then a total protein stain was run according to manufacturer’s procedure (Li-Cor). Following protein quantification, the membrane was incubated in TBS blocking buffer (Li-Cor) for 1 h, and then purified anti-HA mouse monoclonal antibody (Biolegend) in TBS blocking buffer (1:2,500 dilution) for 1 hr. Following primary antibody detection, the membrane was washed 4 times in TBST 0.1%. Secondary detection was done by incubation with goat anti-mouse in TBS blocking buffer (1:15000 dilution) for 1 hour in the dark. Following detection, the membrane was washed 3 times in TBST 0.1% and once in TBS. The membrane was then dried and imaged on the Li-Cor Odyssey CLx imager relative to REVERT total protein stain.

## Acknowledgements

Research reported in this publication was supported by grants from the National Institutes of Health to D.A.H. (R01 GM108492 to D.A.H) and HOGAN19G0, and NSF 1458359 (D.A.H. and D.L.M). Support for D.L.M came in part from NIH/NIAID T32AI007519 (D.L.M). NIGMS P20GM113132 through the Molecular Interactions and Imaging Core (MIIC). STANTO19R0 from the Cystic Fibrosis Foundation and NIDDK P30-DK117469 (Dartmouth Cystic Fibrosis Research Center). RNA-Seq and NanoString were carried out at Dartmouth Medical School in the Genomics Shared Resource, which was established by equipment grants from the NIH and NSF and is supported in part by a Cancer Center Core Grant (P30CA023108) from the National Cancer Institute. We also thank Georgia Doing for preliminary RNAseq analysis, Pat Occhipinti for swapping the antibiotic marker on the *rhlI-*HA expression vector, Carla Cugini for strain construction, and Alan Collins for the gradient plate prop.

## Figure legends

**Figure S1.**
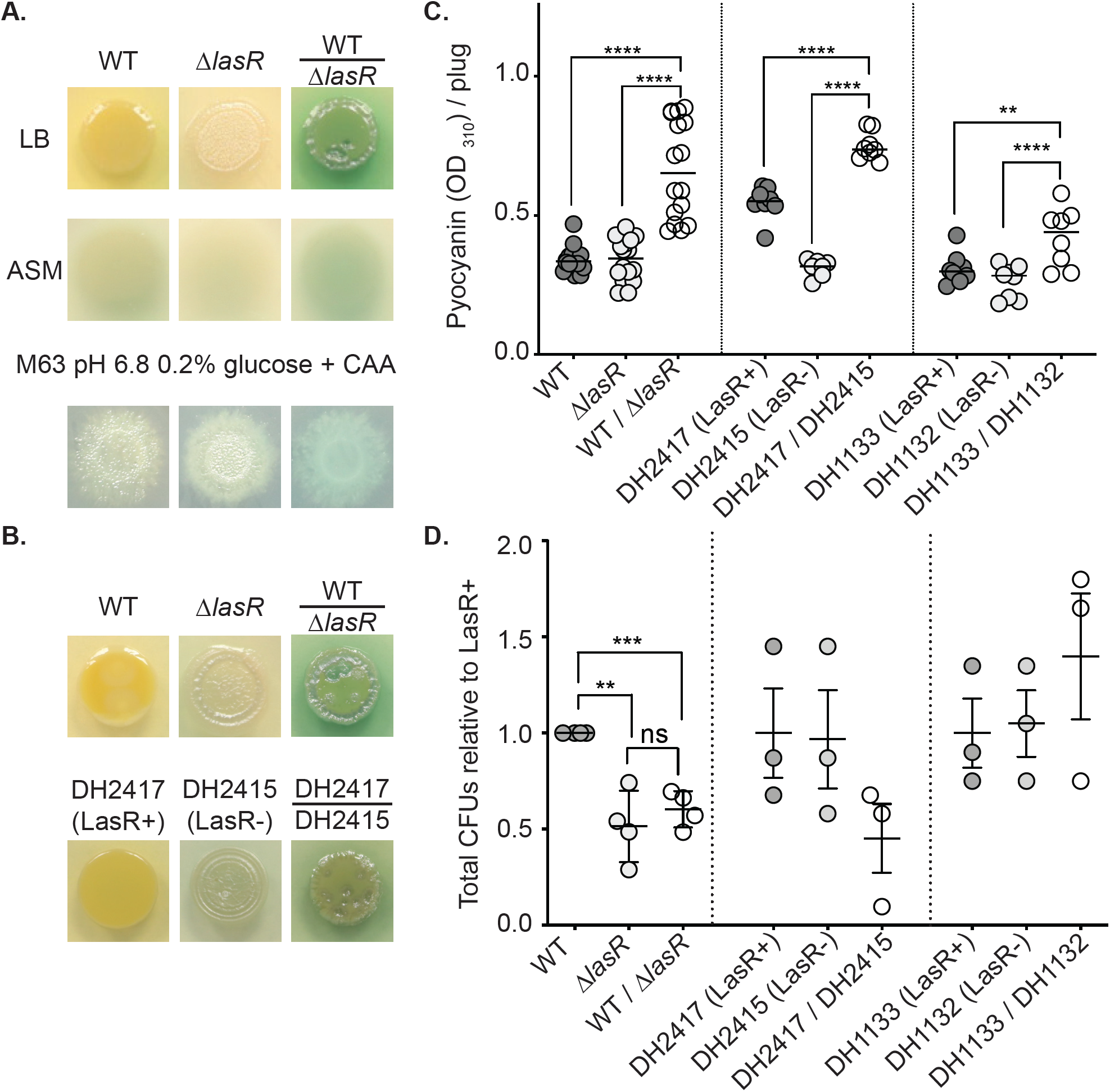
Increased pyocyanin production of LasR+ and LasR- strain co-cultures relative to either strain alone is stable across media and strains. **A**. Representative images from PA14 wild type (WT) and Δ*lasR* mono- and co-culture pyocyanin production on rich media (LB and Artificial Sputum Media-ASM) and buffered minimal medium (M63 pH 6.8 0.2% glucose 0.2% CAA). **B**. Pyocyanin production of mono- and co-cultures of strain PA14 WT and Δ*lasR* and clonally-derived clinical isolates DH2417 (LasR+) and DH2415 (LasR-). **C**. Pyocyanin concentrations of LasR+ and LasR-strains grown in mono- and co-culture on 96 well LB agar plugs for 16 h. Strain PA14 WT and Δ*lasR*, and previously characterized clinical isolate pairs DH2417 (LasR+) and DH2415 (LasR-) and DH1133 (LasR+) and DH1132 (LasR-), from two distinct CF patients, are also shown. Data are from two independent experiments with four biological replicates each. *\*\*\*\** represents a significant difference from the co-culture, p<0.0001. **D**. Colony forming units (CFUs) in 16 h colony biofilms grown on LB. Each strain set is presented relative to WT or the LasR+ strain. PA14 WT and Δ*lasR* mono- and co-cultures data are the average from four independent experiments with at least three biological replicates. No significant (ns) differences found by ANOVA with Tukey multiple hypotheses correction for clinical isolate CFU counts. ** p-value < 0.005, *** p-value < 0.0005.

**Figure S2.**
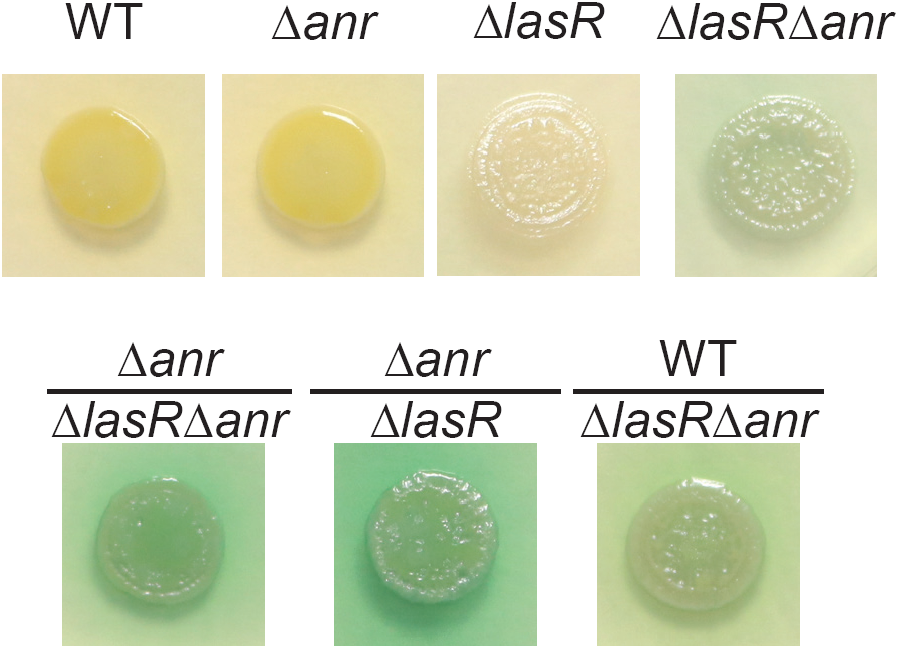
Δ*lasR* shows Anr independent increases in pyocyanin in co-culture with LasR+ *P. aeruginosa*. Pyocyanin production of WT, Δ*anr*, and Δ*lasR*Δ*anr* in mono- and co-culture colony biofilms on LB after 24 h.

**Figure S3.**
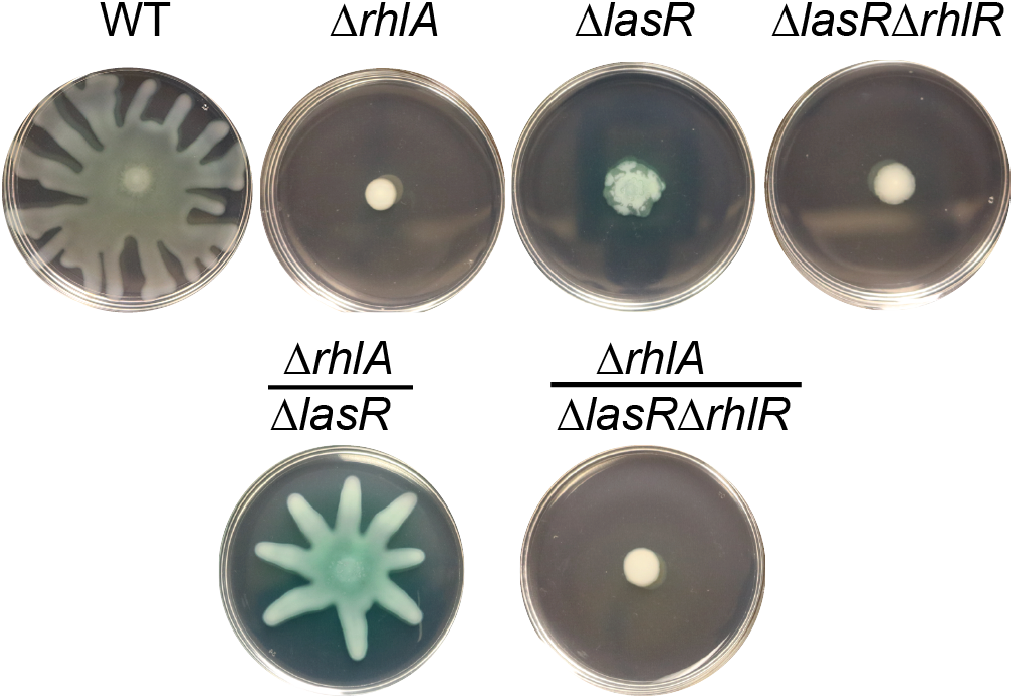
Δ*lasR* shows enhanced production of RhlR-regulated rhamnolipid surfactant in co-culture with LasR+ strain. RhlR-regulated swarming motility on soft agar of WT, rhamnolipid surfactant biosynthesis mutant (Δ*rhlA*), Δ*lasR*, and Δ*lasR*Δ*rhlR* in mono- (top) and co-culture (bottom) after 36 h.

**Figure S4.**
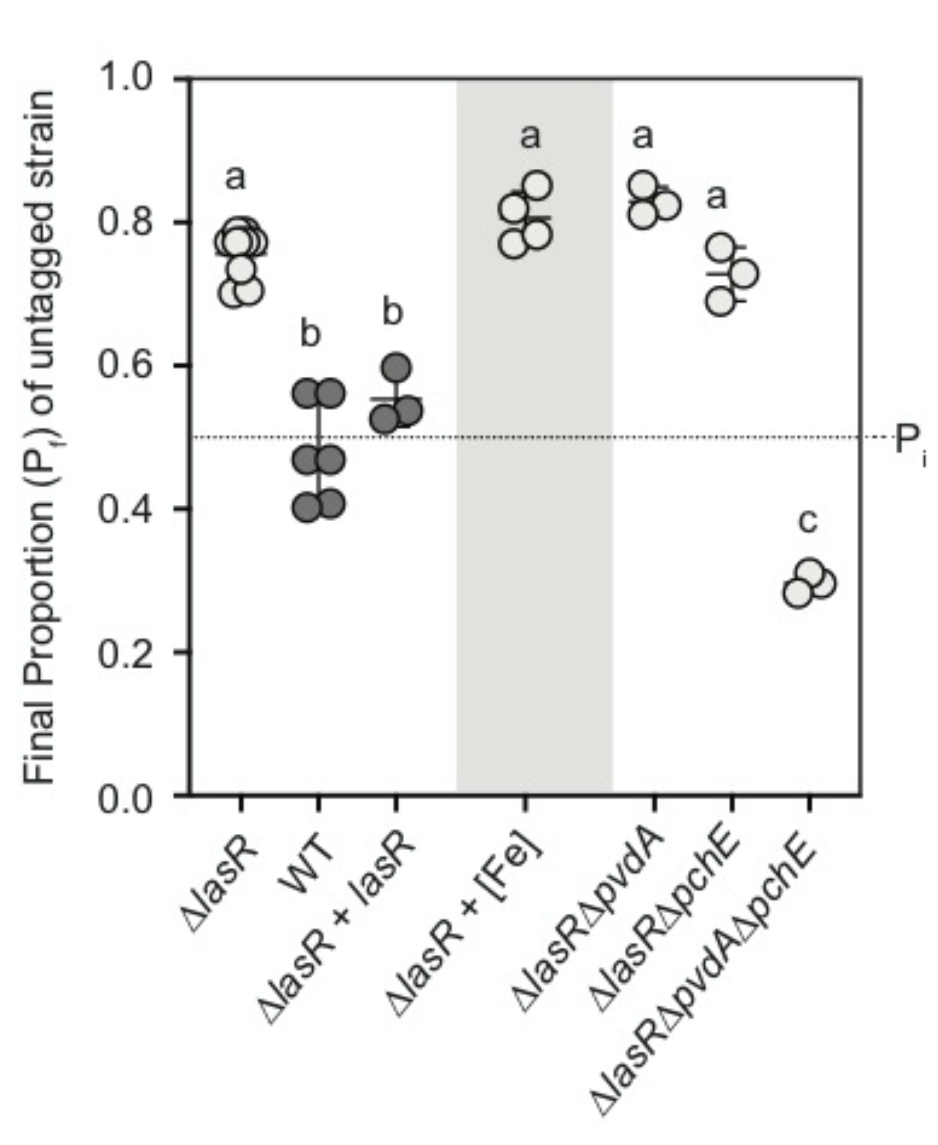
Fitness of Δ*lasR* lacking one or both major siderophore biosynthesis pathways. Final proportion of untagged colony forming units quantified after 16 h competition with *att::lacZ* wild type. Dotted line indicates 0.5 initial proportion (P_i_). Final proportions (P_f_) for Δ*lasR*, wild type (WT), and Δ*lasR* complemented with the *lasR* gene at the native locus (Δ*lasR +* lasR) on LB (white background), for Δ*lasR* on LB supplemented with 10 µM FeSO4 (grey background), and siderophore deficient Δ*lasR* derivatives. Statistical analyses performed by one-way ANOVA for comparison to Δ*lasR* on LB with Sidak multiple hypotheses correction: a-b p-value < 0.0006, a-c p-value < 0.0001.

**Figure S5.**
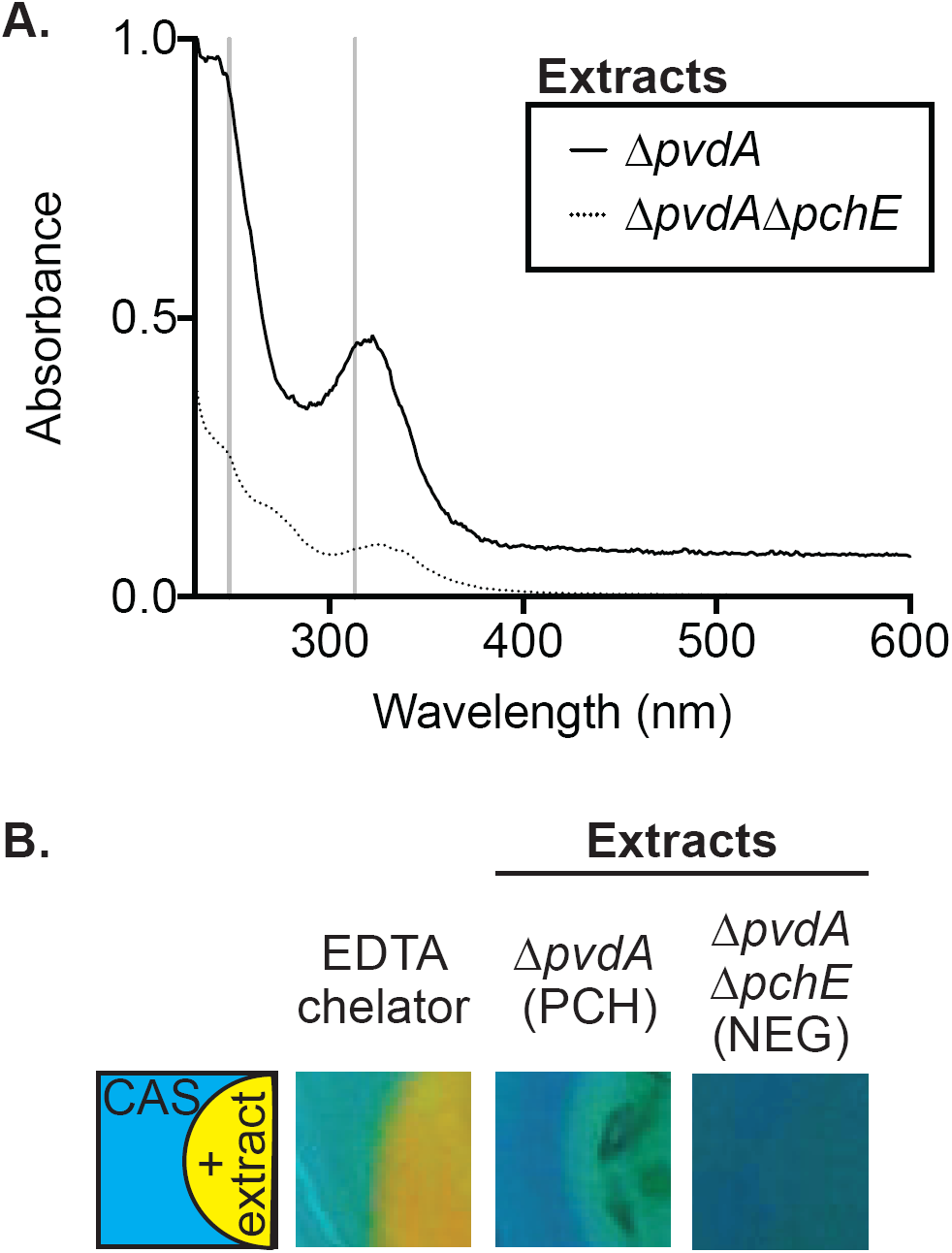
Pyochelin-containing extracts are biologically active. **A**. Absorbance spectra (230 to 600 nm) of extracts from supernatants of pyochelin-producing, Δ*pvdA* (solid line), and pyochelin-deficient, Δ*pvdA*Δ*pchE* (dotted line) strains in a 50/50 MeOH/dH_2_O solution. Grey vertical lines indicate reported peaks for purified, iron-free pyochelin at 248 and 313 nm. **B**. Indicated extracts spotted on chrome azurol S (CAS) agar with ethylenediaminetetraacetic acid (EDTA) metal chelator as positive control wherein change from blue to yellow indicates iron chelating capacity. CAS activity for the extracts from Δ*pvdA* (+PCH) and Δ*pvdA*Δ*pchE* (NEG) are shown.

**Figure S6.**
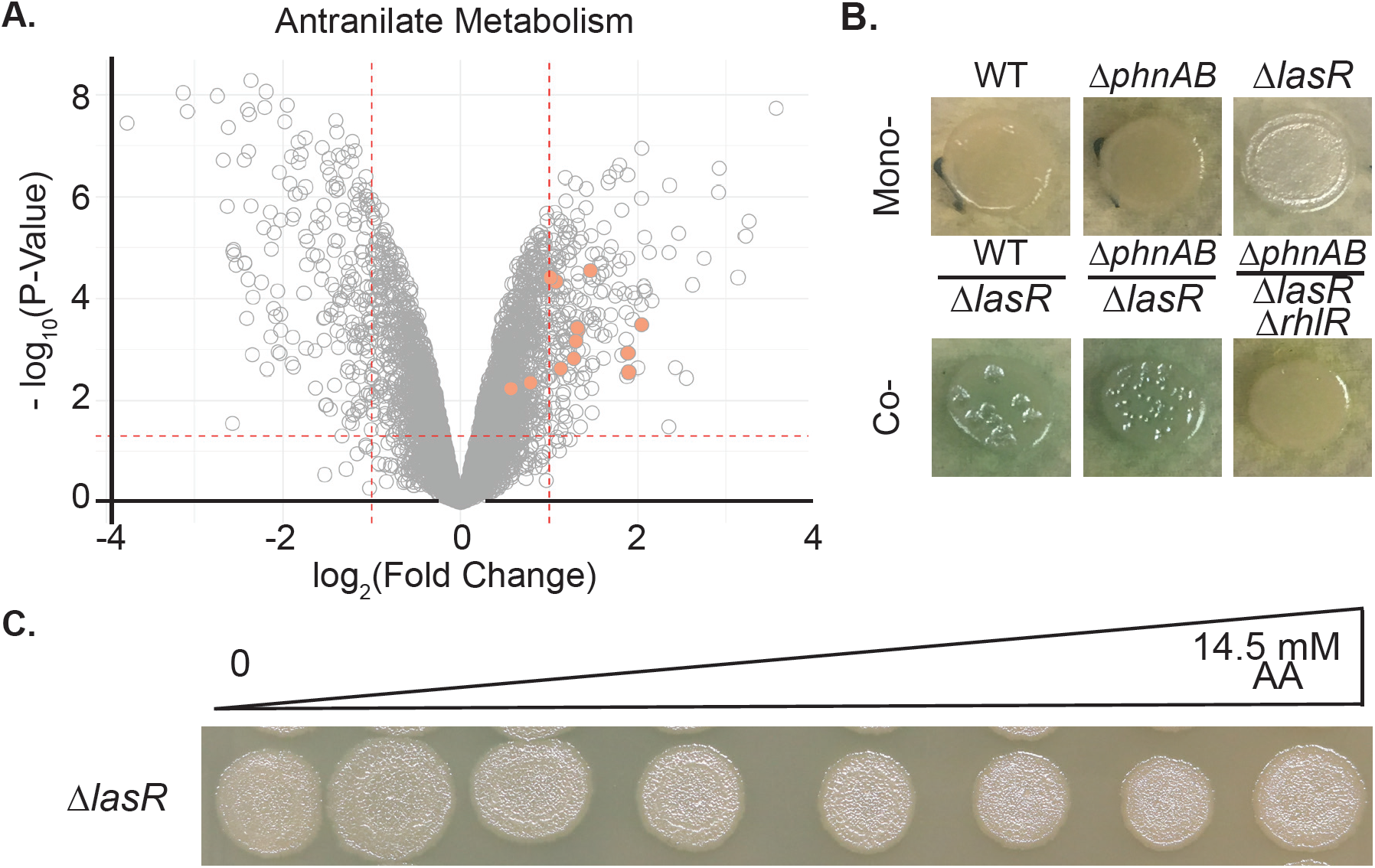
Anthranilate (AA) did not stimulate Δ*lasR* pyocyanin production. **A**. Volcano plot indicating anthranilate metabolism gene expression (orange) in Δ*lasR* grown in co-culture with WT. **B**. RhlR dependent pyocyanin production of Δ*lasR* and anthranilate synthase mutant (Δ*phnAB*) mono- and co-cultures on LB after 18 h. **C**. Colony morphology and pyocyanin production are not different upon anthranilate supplementation across a gradient of concentrations.

**Figure S7.**
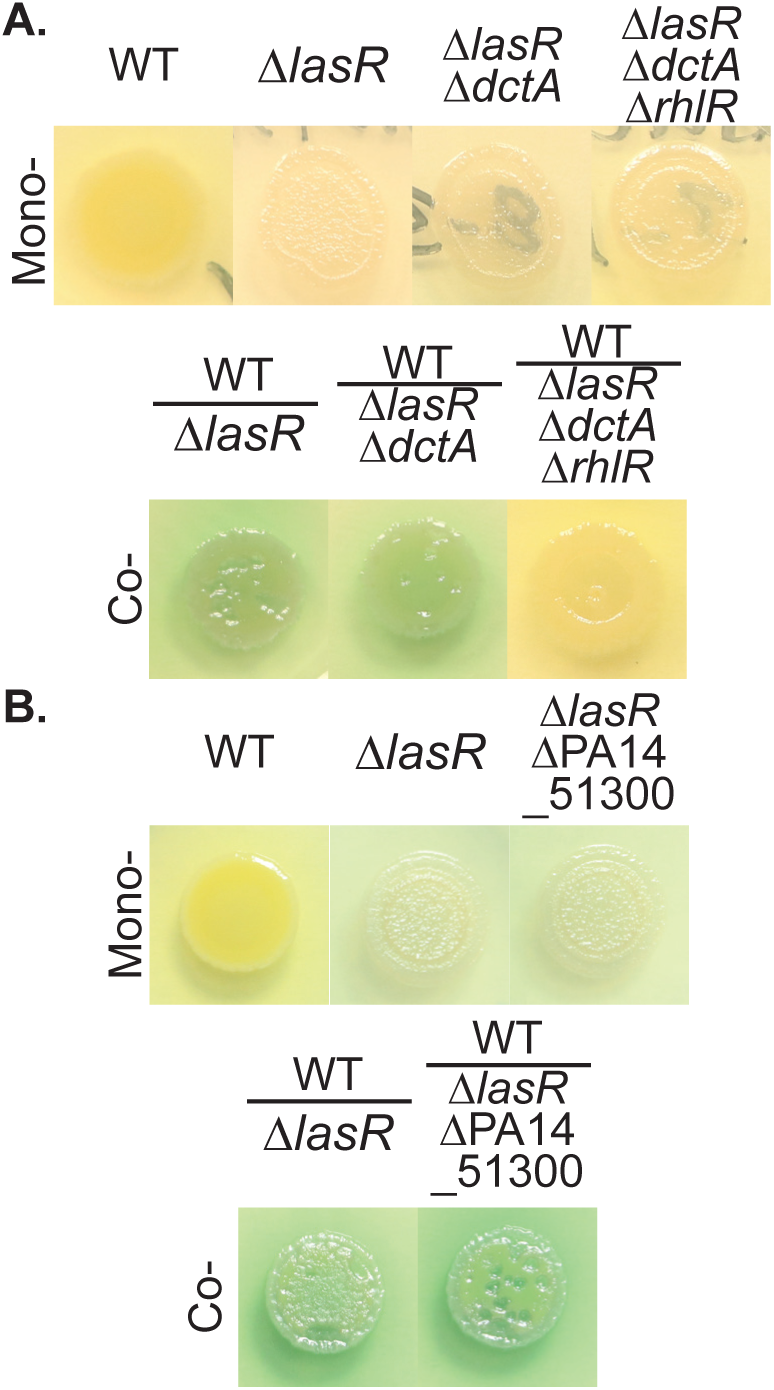
Potential di- and tri-carboxylic acid transporters *dctA* and PA14_51300 were not required for Δ*lasR* pyocyanin production. **A**. Major succinate, fumarate, and malate transporter *dctA* was dispensable in Δ*lasR* background for RhlR-dependent pyocyanin production in co-culture with WT. **B**. Broad TCA cycle intermediate transporter PA14_51300 was not required in Δ*lasR* for pyocyanin production in co-culture with WT.

## Notes

### Competing Interest Statement

The authors have declared no competing interest.

