## Supplemental Table S2 for "Intra-species signaling between *Pseudomonas aeruginosa* genotypes increases production of quorum sensing controlled virulence factors"

**Table S2. Strains, plasmids, and primers used in this study**

| *Strain* | *Strain.ID* | *Description* | *Source* |
|---|---|---|---|
| *P. aeruginosa* | | | |
| PA14 WT | DH122 | Laboratory reference strain | (1) |
| PA14 Δ*lasR* | DH164 | DH122 with in-frame deletion of *lasR* (*PA14_45960*) | (2) |
| PA14 Δ*phz* | DH933 | In-frame deletions of *phzA1-G1* and *phzA2-G2* | (3) |
| PA14 Δ*lasR*Δ*phz* | DH236 | In-frame deletions of *lasR* (*PA14_45960*), *phzC1*, and *phzC2* | This study |
| PA14 WT *att::lacZ* | DH22 | PA14 WT with constitutive expression of *lacZ* | Roberto Kolter (4, 5) |
| PA14 Δ*lasR* + *lasR* | DH3398 | DH164 with complementation of *lasR* (*PA14_45960*) at the native locus | (6) |
| NC-AMT0101-1-2 | DH2417 | Chronic CF lung infection isolate with functional LasR allele, parent of NC-AMT0101-1 | (7) |
| NC-AMT0101-1-1 | DH2415 | Chronic CF lung infection isolate related to DH2417 with LasR LOF (frameshift) allele | (7) |
| AMT0047-2 | DH1133 | Chronic CF clinical isolate with functional LasR allele, parent of AMT0047-3 | (7) |
| AMT0047-3 | DH1132 | Chronic CF clinical isolate related to DH1133 with LasR LOF (premature termination) allele | (7) |
| PA14 Δ*lasR*Δ*anr* | DH2401 | In-frame deletion of *lasR* (*PA14_45960*) and *anr* (*PA14_44490*) | (8) |
| PA14 Δ*anr* | DH2855 | DH122 with in-frame deletion of *anr* (*PA14_44490*) | (9) |

| | | | |
|---|---|---|---|
| PA14 Δ*lasR*Δ*pqsR* | DH1111 | In-frame deletion of *lasR* in DH1110 (Δ*pqsR*) | (10) |
| PA14 Δ*lasR*Δ*rhlR* | DH2944 | In-frame deletion of *lasR* and *rhlR* | (6) |
| PA14 Δ*lasR*Δ*rhlI* | DH238 | In-frame deletion of *lasR* in DH169 | (10) |
| PA14 Δ*lasR* P*rhlI-lacZ* | DH3313 | PA14 Δ*lasR* (DH164) expressing P*rhlI-lacZ* promoter fusion at the *att*::Tn7 site | This study |
| PA14 Δ*lasR*Δ*rhlR* P*rhlI-lacZ* | DH3309 | PA14 Δ*lasR*Δ*rhlR* (DH2944) expressing P*rhlI-lacZ* promoter fusion at the *att*::Tn7 site | (6) |
| PA14 Δ*lasR* P*pqsA-lacZ* | DH3786 | PA14 Δ*lasR* (DH164) expressing P*pqsA-lacZ* promoter fusion at the *att*::Tn7 site | This study |
| PA14 Δ*lasR*Δ*pqsR* P*pqsA-lacZ* | DH3787 | PA14 Δ*lasR*Δ*pqsR* (DH1111) expressing P*pqsA-lacZ* promoter fusion at the *att*::Tn7 site | This study |
| PA14 Δ*pqsA* | DH556 | In-frame deletion of *pqsA* | (11) |
| PA14 Δ*rhlI* | DH169 | In-frame deletion of *rhlI* | (2) |
| PA14 Δ*lasI*Δ*rhlI* | DH242 | In-frame deletions of *lasI* and *rhlI* | This study |
| PA14 Δ*rhlA* | DH7 | Allelic replacement of *rhlA*, $Gm_R$ | (12) |
| PA14 Δ*pvdA* | DH3788 | In-frame deletion of *pvdA* | (13) |
| PA14 Δ*pchE* | DH3789 | In-frame deletion of *pchE* | (13) |
| PA14 Δ*pvdA*Δ*pchE* | DH3790 | In-frame deletions of *pvdA* and *pchE* | (14) |
| PA14 Δ*lasR*Δ*pvdA*Δ*pchE* | DH3791 | In-frame deletion of *lasR* in Δ*pvdA*Δ*pchE* (DH3790) | This study |
| PA14 Δ*lasR*Δ*pvdA* | DH3792 | In-frame deletion of *lasR* in Δ*pvdA* (DH3788) | This study |
| PA14 Δ*lasR*Δ*pchE* | DH3793 | In-frame deletion of *lasR* in Δ*pchE* (DH3789) | This study |

| | | | |
|---|---|---|---|
| PA14 WT P*rhlI-lacZ* | DH3308 | PA14 WT (DH122) expressing P*rhlI-lacZ* promoter fusion at the *att*::Tn7 site | (6) |
| PA14 Δ*lasR* + pmQ72_*rhlI*-HA | DH3798 | PA14 Δ*lasR* (DH164) expressing arabinose-inducible, extrachromosomal HA-tagged rhlI; $Gm_R$ | This study |
| PA14 Δ*lasR* + *lasR* + pmQ72_*rhlI*-HA | DH3799 | PA14 Δ*lasR* + *lasR* (DH3398) expressing arabinose-inducible, extrachromosomal HA-tagged rhlI; $Gm_R$ | This study |
| 388D | DH2606 | SCUT clinical isolate with LasR LOF allele (I215S substitution); MLST type 244 | (15, 16) |
| 550A | DH2615 | SCUT clinical isolate with functional LasR allele; MLST type 244 | (15, 16) |
| 388D + pmQ72_*rhlI*-HA | DH3800 | Clinical isolate 388D (DH2606) expressing arabinose-inducible, extrachromosomal 6x HA-tagged rhlI; $Gm_R$ | This study |
| NC-AMT0101-1-1 + pmQ72_*rhlI*-HA | DH3801 | Clinical isolate NC-AMT0101-1-1 (DH2415) expressing arabinose-inducible, extrachromosomal 6x HA-tagged rhlI; $Gm_R$ | This study |
| 550A + pmQ72_*rhlI*-HA | DH3802 | Clinical isolate 550A (DH2615) expressing arabinose-inducible, extrachromosomal 6x HA-tagged rhlI; $Gm_R$ | This study |
| NC-AMT0101-1-2 + pmQ72_*rhlI*-HA | DH3803 | Clinical isolate NC-AMT0101-1-2 (DH2417) expressing arabinose-inducible, extrachromosomal 6x HA-tagged rhlI; $Gm_R$ | This study |
| PA14 Δ*lasRclpX::*Tn*M* | DH317 | Tn*M* disruption of *clpX* in DH164; $Gm_R$ | This study |
| PA14 Δ*lasRclpX::TnM* + pmQ70_*rhlI*-HA | DH3806 | PA14 Δ*lasRclpX::*Tn*M* (DH317) expressing arabinose-inducible, | This study |

| | | | |
|---|---|---|---|
| | | extrachromosomal 6x HA-tagged rhlI; $Gm_R$, $Amp_R$ | |
| PA14 Δ*lasR* + pmQ70_EV | DH3804 | PA14 Δ*lasR* (DH164) expressing pmQ70 empty expression vector; $Amp_R$ | This study |
| PA14 Δ*lasR* + pmQ70_*rhlI*-HA | DH3805 | PA14 Δ*lasR* (DH164) expressing arabinose-inducible, extrachromosomal 6x HA-tagged rhlI | This study |
| PA14 Δ*lasR* + pmQ72 EV | DH3797 | PA14 Δ*lasR* (DH164) expressing pmQ72 empty expression vector; $Gm_R$ | This study |
| PA14 Δ*rhlI* + pmq72_EV | DH3795 | PA14 Δ*rhlI* (DH169) expressing pmQ72 empty expression vector; $Gm_R$ | This study |
| PA14 Δ*rhlI* + pmq72_*rhlI*-HA | DH3796 | PA14 Δ*rhlI* (DH169) expressing arabinose-inducible, extrachromosomal 6x HA-tagged rhlI; $Gm_R$ | This study |
| PA14 Δ*phnAB* | DH15 | In-frame deletions of *phnA* and *phnB* | (17) |
| PA14 Δ*lasR*Δ*PA14_51300* | DH3807 | In-frame deletion of *PA14_51300* in DH164 | This study |
| PA14 Δ*lasR*Δ*dctA* | DH3811 | In-frame deletion of *dctA* in DH164 | This study |
| PA14 Δ*lasR*Δ*rhlR*Δ*dctA* | DH3812 | In-frame deletion of *dctA* in DH2944 | This study |
| *E. coli* | | | |
| S17 λpir | DH71 | Used as a conjugation partner for introducing pMQ30 and GH121-based plasmids. | |
| pMQ30 EV | DH962 | Allelic replacement vector, $Gm_R$ | (18) |
| pMQ72 EV | DH3733 | Arabinose inducible expression vector, $Gm_R$ | (18) |

| | | | |
|---|---|---|---|
| pMQ70 EV | DH1682 | Arabinose inducible expression vector, Amp$^R$ | (18) |
| GH121 EV | DH2830 | For inserting sequences at the *att*::Tn7 site via allelic replacement; Gm$^R$ | (19) |
| GH121_P*rhlI-lacZ* | DH3314 | lacZ under control of the *rhlI* promoter, for integration at the *att*::Tn7 site; Gm$^R$ | (6) |
| GH121_P*pqsA-lacZ* | DH3785 | lacZ under control of the *pqsA* promoter, for integration at the *att*::Tn7 site; Gm$^R$ | This study |
| p*lasR*_KO | DH133 | PA14 *lasR* in-frame deletion construct; Gm$^R$ | (2) |
| pMQ30_*dctA*_KO | DH3808 | PA14 *dctA* in-frame deletion construct; Gm$^R$ | This study |
| pMQ30_*PA14_51300*_KO | DH3697 | PA14_51300 in-frame deletion construct; Amp$^R$ | This study |
| pMQ72_*rhlI*-HA | DH3794 | Vector for arabinose-inducible gene expression of 6 x HA-tagged RhlI; Gm$^R$ | This study |
| pMQ70_*rhlI*-HA | DH3608 | Vector for arabinose-inducible gene expression of 6 x HA-tagged RhlI; Amp$^R$ | This study |

1. Rahme LG, Stevens EJ, Wolfort SF, Shao J, Tompkins RG, Ausubel FM. 1995. Common virulence factors for bacterial pathogenicity in plants and animals. Science 268:1899-902.
2. Hogan DA, Vik A, Kolter R. 2004. A *Pseudomonas aeruginosa* quorum-sensing molecule influences *Candida albicans* morphology. Mol Microbiol 54:1212-23.
3. Dietrich LE, Price-Whelan A, Petersen A, Whiteley M, Newman DK. 2006. The phenazine pyocyanin is a terminal signalling factor in the quorum sensing network of *Pseudomonas aeruginosa*. Mol Microbiol 61:1308-21.

4. Wang Z, Xiong G, Lutz F. 1995. Site-specific integration of the phage phi CTX genome into the *Pseudomonas aeruginosa* chromosome: characterization of the functional integrase gene located close to and upstream of *attP*. Mol Gen Genet 246:72-9.
5. Choi KH, Schweizer HP. 2006. mini-Tn7 insertion in bacteria with single *att*Tn7 sites: example *Pseudomonas aeruginosa*. Nat Protoc 1:153-61.
6. Harty CE, Martins D, Doing G, Mould DL, Clay ME, Occhipinti P, Nguyen D, Hogan DA. 2019. Ethanol stimulates trehalose production through a SpoT-DksA-AlgU dependent pathway in *Pseudomonas aeruginosa*. Journal of Bacteriology doi:10.1128/jb.00794-18:JB.00794-18.
7. Smith EE, Buckley DG, Wu Z, Saenphimmachak C, Hoffman LR, D'Argenio DA, Miller SI, Ramsey BW, Speert DP, Moskowitz SM, Burns JL, Kaul R, Olson MV. 2006. Genetic adaptation by *Pseudomonas aeruginosa* to the airways of cystic fibrosis patients. Proc Natl Acad Sci U S A 103:8487-92.
8. Hammond JH, Dolben EF, Smith TJ, Bhuju S, Hogan DA. 2015. Links between Anr and quorum sensing in *Pseudomonas aeruginosa* biofilms. J Bacteriol 197:2810-20.
9. Crocker AW, Harty CE, Hammond JH, Willger SD, Salazar P, Botelho NJ, Jacobs NJ, Hogan DA. 2019. *Pseudomonas aeruginosa* ethanol oxidation by AdhA in low-oxygen environments. J Bacteriol 201.
10. Cugini C, Morales DK, Hogan DA. 2010. *Candida albicans*-produced farnesol stimulates *Pseudomonas* quinolone signal production in LasR-defective *Pseudomonas aeruginosa* strains. Microbiology 156:3096-107.
11. Deziel E, Lepine F, Milot S, He J, Mindrinos MN, Tompkins RG, Rahme LG. 2004. Analysis of *Pseudomonas aeruginosa* 4-hydroxy-2-alkylquinolines (HAQs) reveals a role for 4-hydroxy-2-heptylquinoline in cell-to-cell communication. Proc Natl Acad Sci U S A 101:1339-44.
12. Pukatzki S, Kessin RH, Mekalanos JJ. 2002. The human pathogen *Pseudomonas aeruginosa* utilizes conserved virulence pathways to infect the social amoeba *Dictyostelium discoideum*. Proc Natl Acad Sci U S A 99:3159-64.
13. Filkins LM, Graber JA, Olson DG, Dolben EL, Lynd LR, Bhuju S, O'Toole GA. 2015. Coculture of *Staphylococcus aureus* with *Pseudomonas aeruginosa* Drives *S. aureus* towards Fermentative Metabolism and Reduced Viability in a Cystic Fibrosis Model. J Bacteriol 197:2252-64.
14. Wang Y, Wilks JC, Danhorn T, Ramos I, Croal L, Newman DK. 2011. Phenazine-1-Carboxylic Acid Promotes Bacterial Biofilm Development via Ferrous Iron Acquisition. Journal of Bacteriology 193:3606-3617.
15. Srinivasan M, Mascarenhas J, Rajaraman R, Ravindran M, Lalitha P, Glidden DV, Ray KJ, Hong KC, Oldenburg CE, Lee SM, Zegans ME, McLeod SD, Lietman TM, Acharya NR, Steroids for Corneal Ulcers Trial G. 2012. The steroids for corneal ulcers trial: study design and baseline characteristics. Archives of ophthalmology (Chicago, Ill : 1960) 130:151-157.
16. Hammond JH, Hebert WP, Naimie A, Ray K, Van Gelder RD, DiGiandomenico A, Lalitha P, Srinivasan M, Acharya NR, Lietman T, Hogan DA, Zegans ME. 2016. Environmentally endemic *Pseudomonas aeruginosa* strains with mutations in

lasR are associated with increased disease severity in corneal ulcers. mSphere 1.
17. Mahajan-Miklos S, Tan MW, Rahme LG, Ausubel FM. 1999. Molecular mechanisms of bacterial virulence elucidated using a *Pseudomonas aeruginosa-Caenorhabditis elegans* pathogenesis model. Cell 96:47-56.
18. Shanks RM, Caiazza NC, Hinsa SM, Toutain CM, O'Toole GA. 2006. *Saccharomyces cerevisiae*-based molecular tool kit for manipulation of genes from gram-negative bacteria. Appl Environ Microbiol 72:5027-36.
19. Heussler GE, Cady KC, Koeppen K, Bhuju S, Stanton BA, O’Toole GA. 2015. Clustered Regularly Interspaced Short Palindromic Repeat-Dependent, Biofilm-Specific Death of *Pseudomonas aeruginosa* Mediated by Increased Expression of Phage-Related Genes. mBio 6:e00129-15.
